# Euchromatin Peripheral Organization Follows Anterograde Signalling Under Anaesthetic Stress

**DOI:** 10.64898/2026.08.28.747873

**Authors:** Shilpa Chandra, Sakshi Chouhan, Laxmidhar Behera, Chayan Kanti Nandi

## Abstract

Anterograde and retrograde signalling establish bidirectional communication between the nucleus and chloroplasts. Retrograde signals from chloroplasts regulate nuclear gene expression while anterograde signals from the nucleus coordinate chloroplast development and maintain cellular homeostasis. How this bidirectional signalling framework extends beyond locus-specific regulation to shape the global spatial organization of nuclear chromatin across tissues remains unclear. Although anaesthesia can alter chromatin organisation, the role of chloroplast dysfunction in these changes remains unclear. Here, we investigate how chloroplast dysfunction and anaesthesia influence euchromatin and heterochromatin organisation in *Solanum lycopersicum* seedlings across tissues with contrasting photosynthetic competence. Using confocal and super-resolution radial fluctuation (SRRF) imaging with quantitative multiparameter analysis, we identify distinct, tissue-specific chromatin responses to chloroplast disruption and anaesthesia. Notably, anaesthesia induces distinct spatial chromatin changes across tissues that are independent of chloroplast dysfunction, suggesting a direct nuclear response to anaesthesia rather than a chloroplast-mediated retrograde effect. These findings highlight chromatin topology as a potential quantitative biomarker of cellular disruption and provide a framework for investigating anterograde chloroplast–nucleus coordination and stress-responsive nuclear organisation in plants.

## 1. Introduction

Nuclear architecture is a fundamental determinant of gene expression, cellular identity, and organismal development.^1^ In plants, chromatin organisation within the nucleus is defined by the spatial segregation of transcriptionally active euchromatin, marked by histone H3 lysine 4 trimethylation (H3K4me3), from silent heterochromatin, marked by H3K9 trimethylation (H3K9me3) and concentrated at the nuclear periphery as discrete chromocenters.^2–4^ The inner nuclear membrane is scaffolded by plant-specific CROWDED NUCLEI (CRWN) proteins that anchor heterochromatic domains in lamin-associated domains (LADs), directly coupling chromatin positioning to nuclear shape.^5,6^ Chloroplasts communicate with the nucleus through bidirectional signalling. Anterograde signals, governed by HY5 and the PIFs, direct chloroplast differentiation in response to light.^7,8^ In the opposite direction, plastid-to-nucleus retrograde signalling relays continuous information about chloroplast developmental and metabolic status through molecular intermediaries including reactive oxygen species (ROS), 3 -phosphoadenosine 5 - phosphate (PAP), methylerythritol cyclodiphosphate (MEcPP), and tetrapyrrole-related signals which converge on the GUN1 regulatory hub to modulate nuclear gene expression and epigenetic programmes.^9–11^ Recent work demonstrated that these retrograde signals govern a sequential histone modification switch at photosynthesis-associated nuclear gene (PhANG) loci, converting repressive H3K27 trimethylation to activating H3K27 acetylation in a GUN1-dependent, lincomycin-sensitive manner, establishing that chloroplast function directly shapes the nuclear epigenetic landscape.^10,12^ Separately, plants exposed to anaesthetic agents exhibit reversible suppression of organ movement, electrical signalling, and endocytic vesicle recycling, with documented effects on nuclear positioning and chromatin redistribution, pointing to a cytoskeletal contribution to nuclear organisation.^4,13^ Nuclear architecture in plants therefore reflects the integrated state of both organellar signalling and cytoskeletal integrity. ^14^

Despite these advances, two critical questions remain unresolved. First, whether retrograde signalling governs global chromatin spatial topology, the spatial positioning of heterochromatin and euchromatin domains across the nuclear volume rather than merely locus-specific histone modifications, has not been directly tested. Second, whether anaesthesia-induced chromatin reorganisation depends on chloroplast dysfunction as an intermediate, or engages a plastid-independent nuclear pathway, has not been resolved. Answering these questions requires systematic comparison across conditions that selectively disrupt chloroplast function, selectively apply anaesthesia, or combine both perturbations, across tissues that differ in chloroplast content and therefore in the retrograde signalling they receive. This comparison has not previously been performed, representing a significant gap in understanding how organellar and external perturbations converge on nuclear chromatin organisation.

Here, we address this gap using confocal and super-resolution radial fluctuation (SRRF) imaging to characterise chloroplast-nucleus organisation in *Solanum lycopersicum* seedlings across six experimental conditions and three tissue types. To our knowledge, this is the first study to simultaneously characterise the spatial organisation of euchromatin and heterochromatin across leaf, stem, and root tissues in a crop plant subjected to chloroplast perturbation and anaesthetic exposure. We combine macroscopic and pigmentation phenotyping, UV-visible spectrophotometry, chloroplast imaging, and quantitative analysis of chloroplast-nucleus spatial association with SRRF-based mapping of euchromatin (H3K4me3) and heterochromatin (H3K9me3). By examining leaf, stem, and root tissues under chloroplast disruption and anaesthetic exposure, the study further evaluates whether anaesthesia-associated chromatin reorganisation follows patterns linked to chloroplast functional status or represents a distinct, tissue-dependent nuclear response. The findings have broad implications for plant biology and crop science. Establishing that retrograde signalling controls chromatin spatial topology positions nuclear architecture as a downstream indicator of photosynthetic status relevant to stress acclimation. Demonstrating chromatin reorganisation independent of chloroplast status opens new understanding of how different stress like temperature extremes, osmotic shock, and pathogen attack translate into nuclear architectural changes. The quantitative framework and experimental design developed here provide a toolkit for future investigations of chromatin positioning in diverse plant species and stress conditions.

## 2. Results and Discussion

We first characterised the macroscopic and pigment-level responses of *Solanum lycopersicum* seedlings under six experimental conditions using phenotypic assessment, UV-visible spectrophotometry, and confocal imaging of chloroplast-nucleus organisation. We then examined nuclear chromatin architecture in leaf, stem, and root tissues using SRRF microscopy, separately analysing euchromatin (H3K4me3) and heterochromatin (H3K9me3). Their spatial organisation was quantified using multiple topological parameters, including domain number and area, peripheral enrichment, condensation, radial position to identify tissue- and condition-specific patterns of chromatin reorganisation.

### 2.1 Experimental Conditions Produce Distinct Whole-Plant Phenotypes

The six experimental conditions generated distinguishable macroscopic growth phenotypes, confirming effective physiological perturbation (**Figure 1I**). Control-Light plants exhibited the most developed green shoot morphology and served as the reference state. Dark-grown and lincomycin-treated plants showed visible reductions in pigmentation (**Figure 1I**, **Figure S1**), consistent with disrupted chloroplast development.^12,15^ UV-visible absorption profiles confirmed condition-dependent reductions in chlorophyll-associated absorbance (**Figure S1**).^15,16^

**Figure 1.**
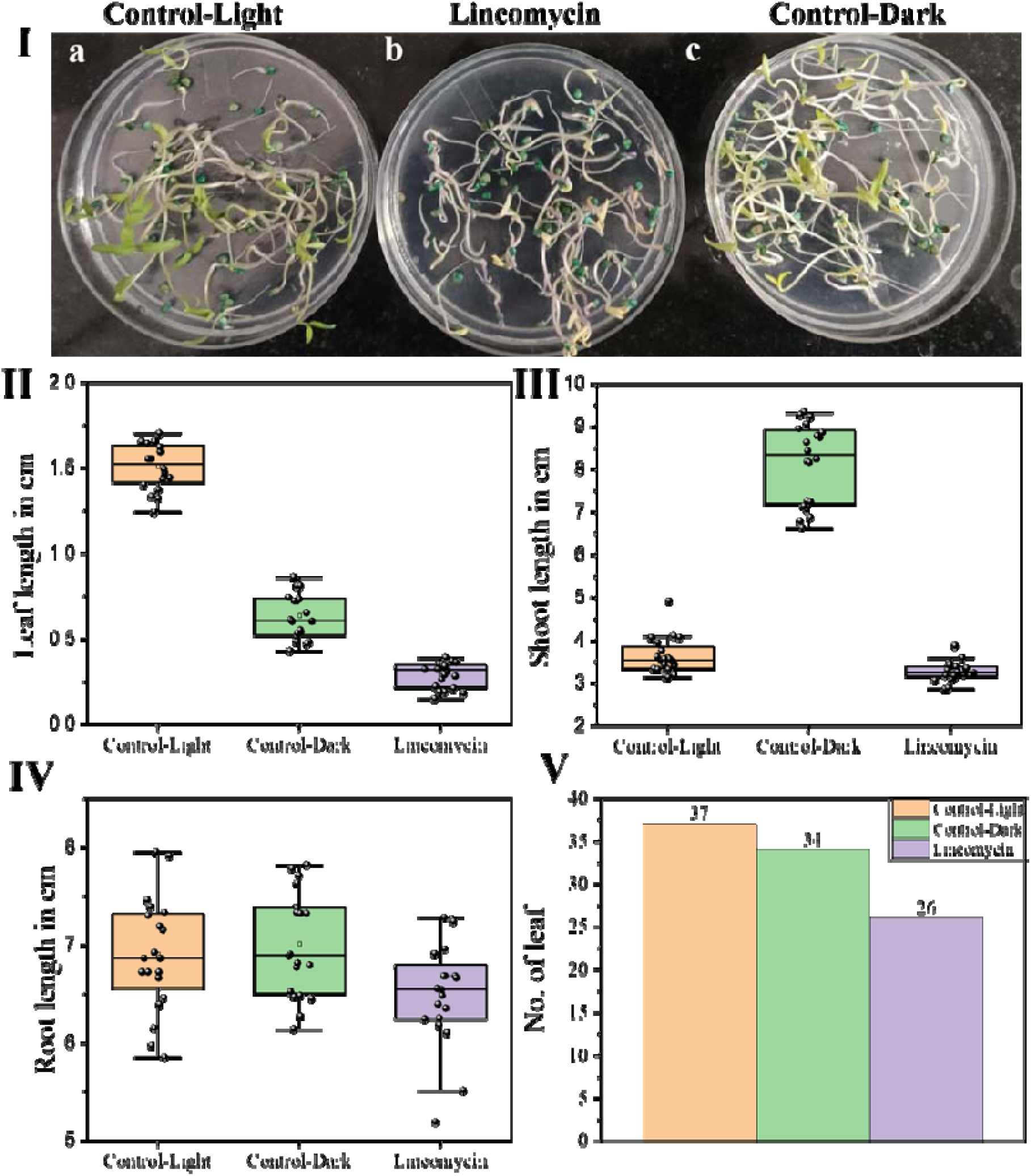
Phenotypic effects of the experimental conditions on plant growth and development. **(I)** Representative images of plants grown under the respective experimental conditions, illustrating differences in overall growth and morphology. **(II)** Quantitative comparison of leaf length (cm) among the experimental groups. **(III)** Quantitative comparison of stem length (cm) among the experimental groups. **(IV)** Quantitative comparison of root length (cm) among the experimental groups. **(V)** Quantitative comparison of leaf number among the experimental groups. Individual points represent measurements from individual plants, with box plots showing the distribution of the observations (n=20).

Organ-level measurements revealed that treatment effects extended systemically (**Figure 1II-V**, n = 20 per condition). Leaf length (**Figure 1II**), stem length **Figure 1III**), root length (**Figure 1IV**), and leaf number (**Figure 1V**) all varied among conditions. Root growth was affected despite the absence of photosynthetically active chloroplasts in root tissue, pointing to systemic effects mediated through the shoot-to-root communication axis.[30] Combined treatments produced phenotypes that were not simple superpositions of individual treatment effects, indicating non-linear physiological interactions at the whole-plant level.^15,17^

### 2.2 Chloroplast-Nucleus Association Is Condition- and Tissue-Dependent

Confocal imaging of chlorophyll autofluorescence and nuclear DAPI fluorescence revealed condition-dependent alterations in chloroplast abundance, morphology, and spatial association with the nucleus in leaf and stem tissues (**Figures 2-3**, **Figures S2-S15**). In Control-Light leaf tissue, well-developed chloroplasts were closely opposed to the nuclear envelope, forming a dense perinuclear shell confirmed by three-dimensional reconstruction (**Figure 2I(a)**, **Figures S2, S4**).^18^ This tight chloroplast-nucleus physical association is consistent with facilitated retrograde signal delivery at the nuclear surface. Chloroplast number and mean area under this condition provided the quantitative baseline (**Figure 2II-III**).^10,12,19^

**Figure 2.**
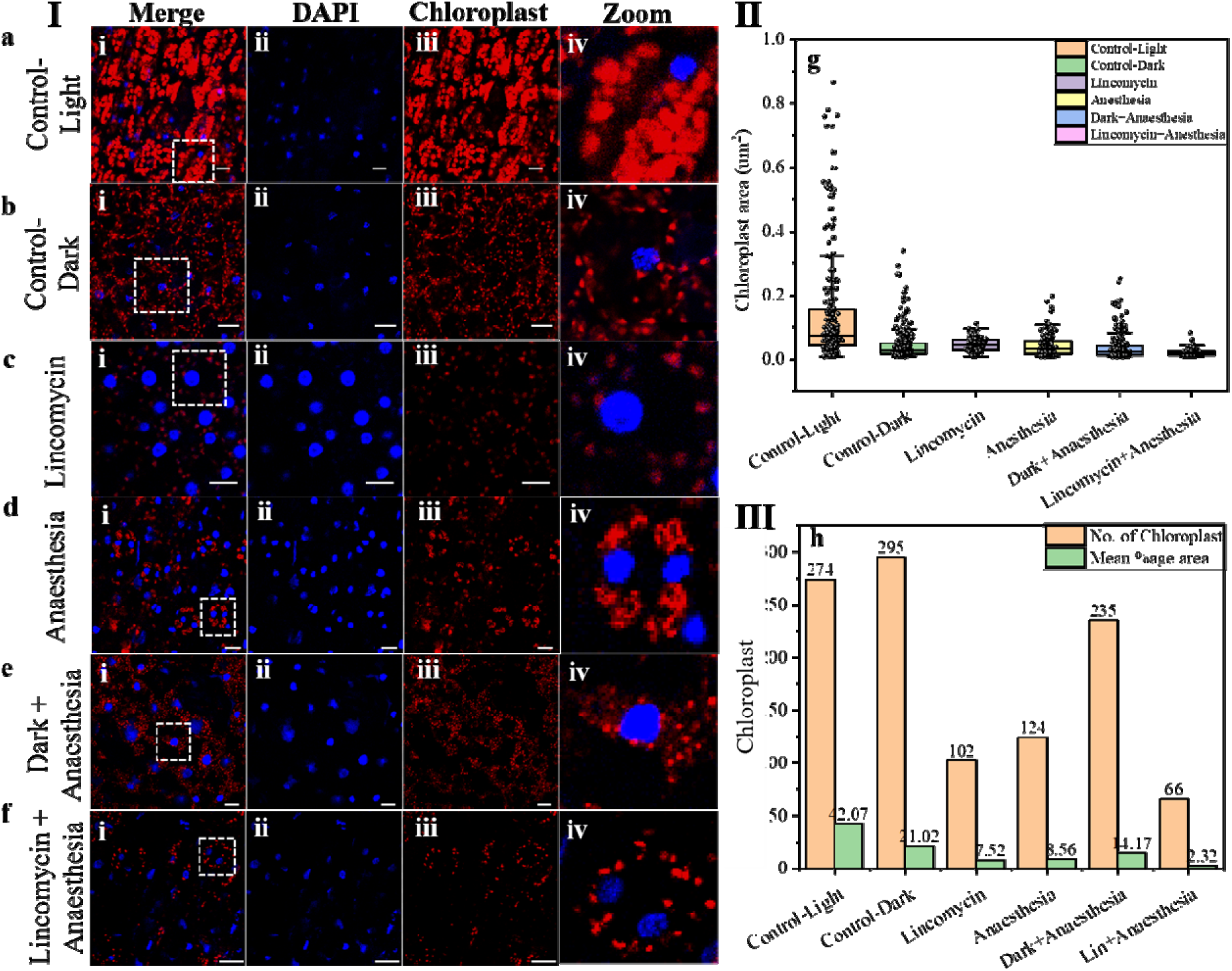
Chloroplast-nucleus association in leaf cells under different experimental conditions. **I(a-f)** Representative fluorescence images showing chloroplasts (red) and nuclear DNA (blue) under the control and experimental conditions. **(a)** Control-light leaf cells show well-developed and regularly distributed chloroplasts, with nuclear DNA distributed throughout the nucleus. **(b-f)** Representative images from the experimental conditions showing condition-dependent alterations in chloroplast size, number, and spatial organization, with comparatively little change in the distribution of nuclear DNA. **II** Quantitative assessment of chloroplast-associated measurements across the experimental conditions. **III** Quantification of chloroplast number and mean chloroplast area across the experimental conditions. n = 10, scale bar = 10 µm.

**Figure 3.**
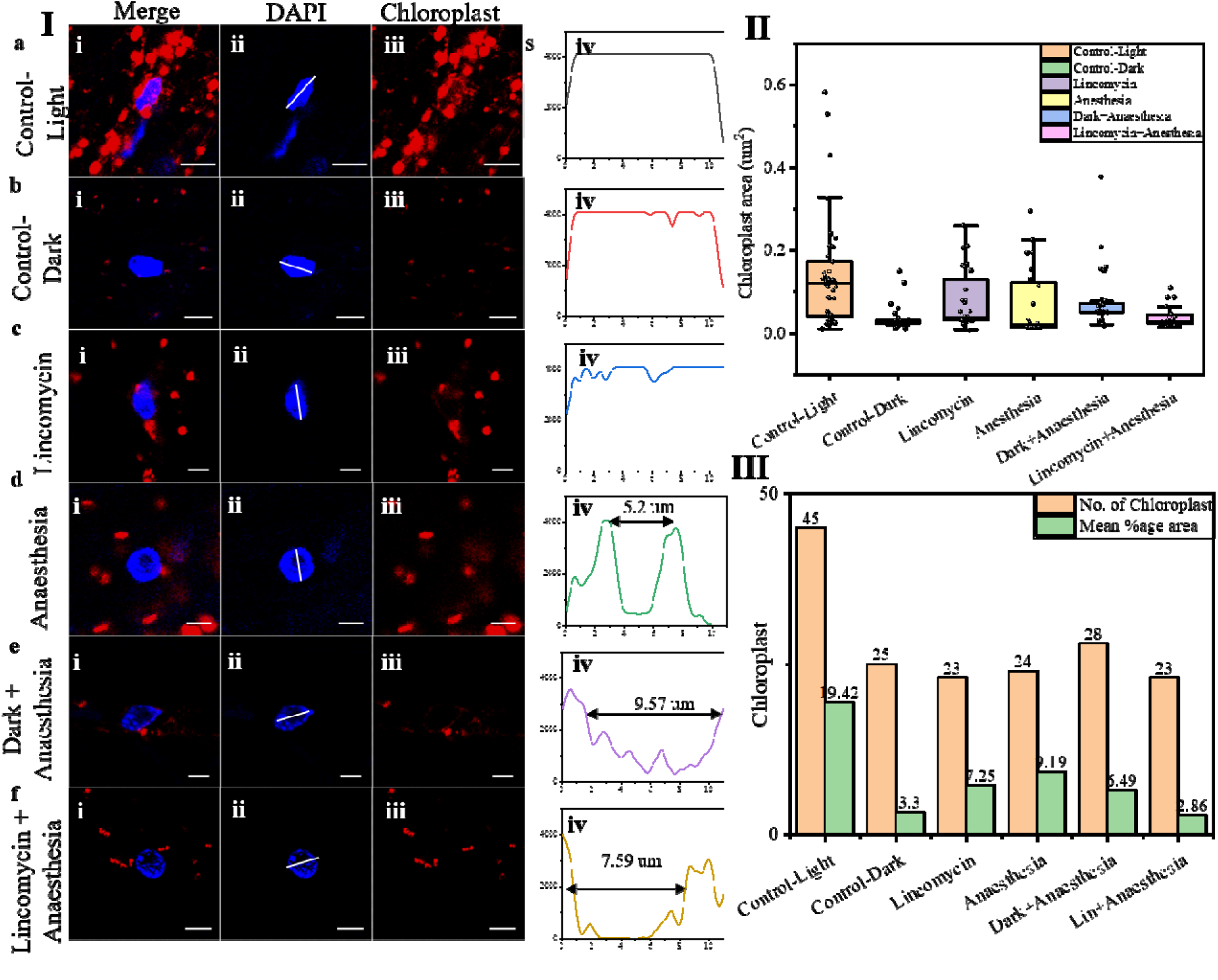
Chloroplast alterations and nuclear DNA redistribution in stem tissue. **I(a-f)** Representative fluorescence images under the respective experimental conditions, showing **(i)** merged chloroplast and nuclear DNA signals, **(ii)** nuclear DNA, **(iii)** chloroplasts, and **(iv)** radial fluorescence intensity profiles of nuclear DNA used to characterize the spatial distribution of DNA within the nucleus. **(a)** Control condition showing well-developed chloroplasts in close association with the nucleus and a relatively distributed nuclear DNA pattern. **(b-c)** Experimental conditions showing reduced chloroplast number and size, with comparatively limited changes i nuclear DNA organization. **(d-f)** Experimental conditions showing further reduction in chloroplast number and size, accompanied by redistribution of nuclear DNA toward the nuclear periphery. **II** Quantitative assessment of chloroplast-associated measurements across the experimental conditions. **III** Quantification of chloroplast number and mean chloroplast area (%) across the experimental conditions. n = 10; scale bar = 10 µm.

Dark-grown seedlings contained etioplast-like structures with reduced autofluorescence and disrupted periorganellar organisation (**Figure 2I(b)**, **Figure S5**).^20,21^ Lincomycin caused pronounced reduction in chloroplast number and fluorescence intensity (**Figure 2I(c)**, **Figure S6**, **Figure 2II-III**), confirming pharmacological inhibition of plastid translation and consequent failure of chloroplast biogenesis.^22,23^ Anaesthesia-containing conditions produced chloroplast fragmentation and reduced periorganellar fluorescence (**Figure 2I(d-f)**, **Figures S7-S9**), indicating a direct physical effect of the anaesthetic on plastid membrane integrity and chloroplast movement.^4,24^

In stem tissue, the baseline chloroplast-nucleus association under Control-Light was moderate (**Figure 3I(a)**, **Figure S3**, **Figure S10**).^25^ Dark and lincomycin conditions reduced chloroplast abundance (**Figure 3I(b-c)**, **Figures S11-S12**, **Figure 3II-III**).^21,25,26^ Anaesthesia-containing conditions produced visible redistribution of nuclear DNA toward the nuclear periphery detectable in the merged fluorescence channel (**Figure 3I(d-f)**, **Figures S13-S15**, **Figure 3I(iv)**), foreshadowing the tissue-specific chromatin responses characterised below. Root tissue, lacking chloroplasts entirely, displayed condition-dependent nuclear DNA distribution changes in response to anaesthesia-containing but not lincomycin-only conditions (**Figure 4I(a-f)**, **Figures S16-S21**, **Figure 4II(a-f)**), establishing that nuclear reorganisation in this tissue cannot be attributed to direct chloroplast perturbation. ^4,27,28^

**Figure 4.**
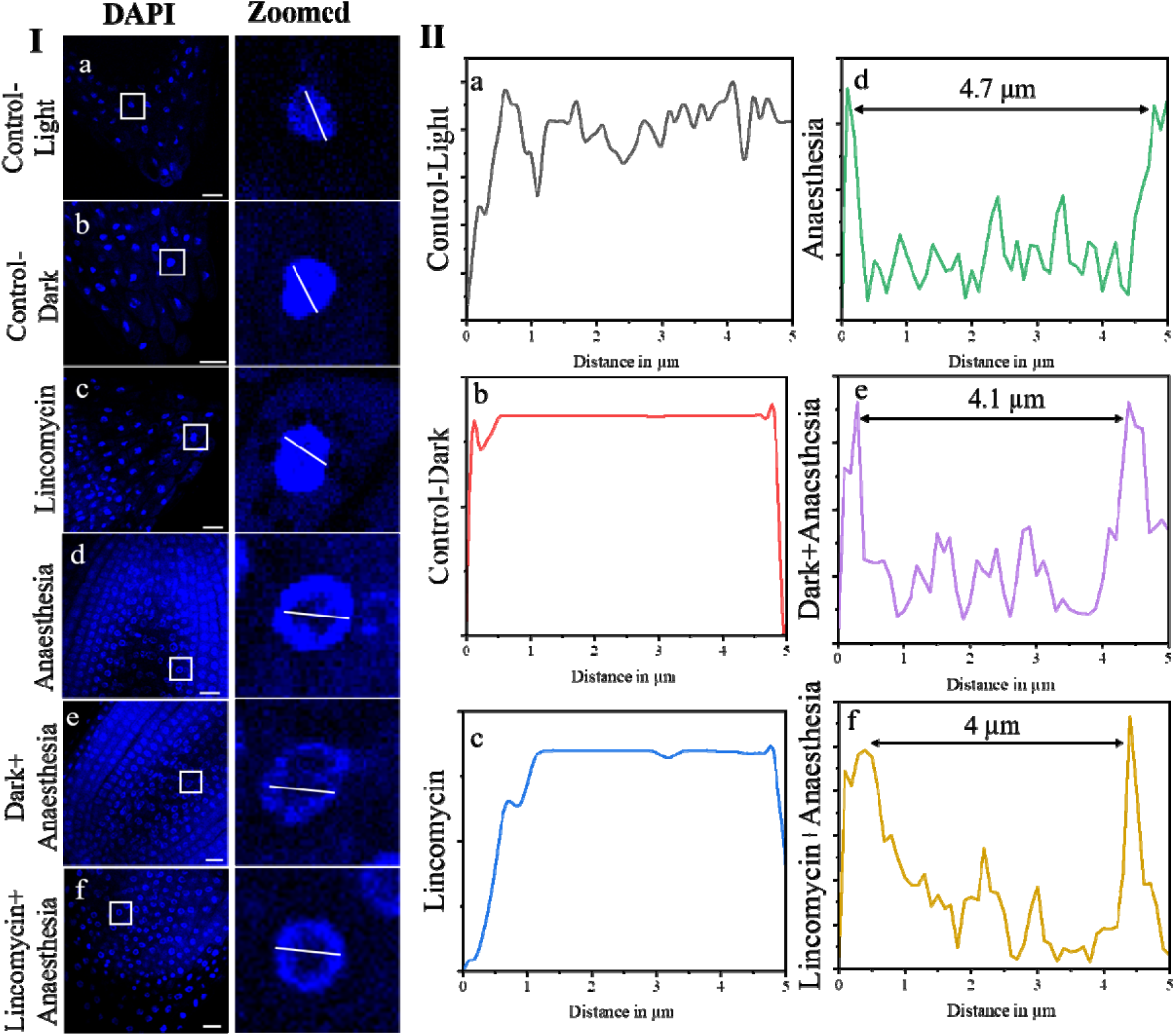
Radial distribution of nuclear DNA in root tissue. Representative fluorescence images of nuclear DNA under the respective experimental conditions are shown in **I (a-f)**. For each condition, the left image shows the overall nuclear DNA distribution, while the adjacent magnified image shows the nucleus with the white line indicating the position of the radial fluorescence intensity scan. **II (a-f)** The corresponding intensity profiles show the spatial distribution of nuclear DNA fluorescence along the indicated transects. The profiles demonstrate condition-dependent changes in nuclear DNA organization, including differences in the relative distribution of DNA between the nuclear periphery and interior. Scale bar = 20 µm.

This systematic characterization of chloroplast status across tissues and conditions provided the physiological context for interpreting nuclear chromatin responses: leaf tissue experienced high retrograde signal input, stem received intermediate input, and root received none. We next examined how these gradients of chloroplast status correlated with nuclear chromatin organization.

### 2.3 Euchromatin Undergoes Condition-Dependent Spatial Transitions

Euchromatin organisation, assessed using H3K4me3 immunofluorescence and SRRF imaging across 18 condition-tissue combinations, demonstrated pronounced condition- and tissue-dependent spatial transitions (**Figure 5**).^29,30^ H3K4me3 is generally associated with transcriptionally active chromatin, and its spatial organisation may therefore reflect the transcriptional and developmental state of individual tissues. Under Control-Light conditions, leaf nuclei exhibited a relatively condensed and spatially restricted euchromatin organisation, whereas stem and root nuclei showed a more dispersed distribution (**Figure 5a**).^4,31,32^ The compact organisation observed in illuminated leaf tissue is consistent with the active transcriptional state associated with photosynthetically competent cells. Chloroplast-derived retrograde signalling can influence nuclear gene expression and chromatin-associated regulation, including transcriptional responses involving photosynthesis-associated nuclear genes (PhANGs).^33^ Under Control-Dark conditions, euchromatin appeared more scattered, particularly in leaf tissue (**Figure 5b**). This change is consistent with reduced photosynthetic activity and altered chloroplast-to-nucleus signalling under darkness. However, because the present study did not directly measure individual retrograde signalling molecules, the observed spatial redistribution should be interpreted as an association with altered chloroplast functional status rather than direct evidence of a specific signalling pathway. ^12,34^

**Figure 5.**
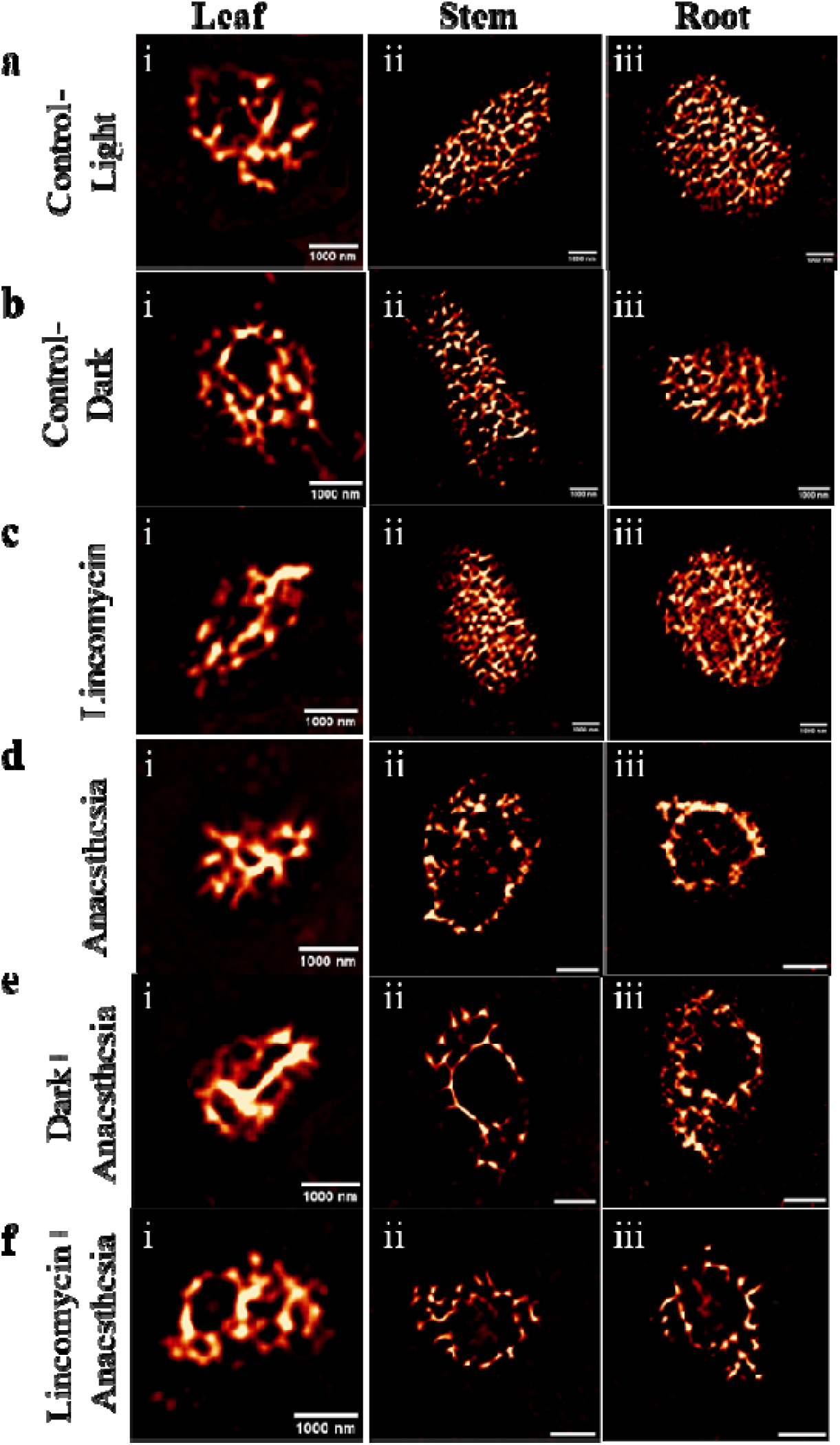
Representative SRRF images showing the spatial organisation of H3K4me3-marked euchromatin across experimental conditions and tissues in *Solanum lycopersicum* seedlings. (**a-f**) correspond to **(a)** Control-Light, **(b)** Control-Dark, **(c)** Lincomycin, **(d)** Anaesthesia, **(e)** Dark + Anaesthesia, and **(f)** Lincomycin + Anaesthesia, respectively. Within each condition, **(i)** leaf, **(ii)** stem, and **(iii)** root nuclei are shown. Under Control-Light and Control-Dark, euchromatin showed a predominantly scattered organisation across all three tissues. Following lincomycin treatment, euchromatin remained scattered in stem and root but appeared more condensed in leaf. In contrast, anaesthesia produced a condensed euchromatin organisation in leaf and a pronounced peripheral arrangement in stem and root. The same tissue-specific pattern was retained under Dark + Anaesthesia an Lincomycin + Anaesthesia, with condensed euchromatin in leaf and peripheral organisation in stem and root. These observations demonstrate a clear tissue-dependent redistribution of euchromatin toward the nuclear periphery under anaesthesia and combined treatments. Scale bars: 1000 nm.

Lincomycin produced a distinct euchromatin phenotype in leaf tissue. Following inhibition of plastid translation, leaf euchromatin appeared more condensed relative to the scattered organisation observed under Control-Dark conditions, while stem and root maintained comparatively dispersed patterns (**Figure 5c)**.^10,12,35^ Lincomycin is widely used to perturb chloroplast function and has been shown to influence nuclear gene expression through disruption of plastid-dependent signalling. The tissue-specific response observed here indicates that disruption of chloroplast translation does not produce a uniform euchromatin phenotype throughout the seedling. The stronger response in leaf may reflect its greater dependence on functional chloroplasts and its more pronounced chloroplast-associated nuclear regulatory environment.^35^

Anaesthesia produced the most distinctive redistribution of euchromatin. In leaf nuclei, H3K4me3-marked euchromatin remained predominantly condensed, whereas stem and root nuclei showed a pronounced shift toward the nuclear periphery (**Figure 5d**). This peripheral organisation was also evident under Dark + Anaesthesia and Lincomycin + Anaesthesia conditions (**Figure 5e, f**), indicating that the effect persisted when chloroplast activity or plastid translation was simultaneously perturbed. This differential response indicates that the anaesthesia-associated redistribution of euchromatin is influenced by the cellular state and is altered when photosynthetic activity or plastid translation is simultaneously perturbed. Thus, the peripheral organisation observed under anaesthesia represents a distinct nuclear response that is not maintained under these combined treatment conditions.^4,36,37^

The quantitative analysis further supported these spatial observations (**Figure 6**). Nuclear area varied across tissues and experimental conditions, demonstrating treatment-dependent changes in nuclear morphology (**Figure 6a**). Changes in euchromatin domain number were particularly evident under anaesthesia, with reductions observed in several stem and root conditions, indicating reorganisation of discrete H3K4me3-marked domains (**Figure 6b**). Corresponding changes in mean domain area suggested that reductions in domain number were accompanied by restructuring or enlargement of individual domains under some conditions (**Figure 6c**). These changes indicate that the treatments altered not only the overall localisation of euchromatin but also its internal domain architecture. Peripheral enrichment provided further evidence for the anaesthesia-associated redistribution. The Peripheral Enrichment Index increased under anaesthesia-containing conditions, particularly in stem and root, consistent with the peripheral patterns observed directly in SRRF images (**Figure 6d**). The increase was retained under both dark + anaesthesia and lincomycin + anaesthesia, suggesting that peripheral euchromatin organisation in these tissues persists despite simultaneous perturbation of chloroplast function. Euchromatin condensation also varied considerably across conditions and tissues (**Figure 6e**), with reduced condensation under several anaesthesia-associated conditions. This indicates that peripheral localisation does not necessarily correspond to increased global chromatin compaction; euchromatin can become spatially peripheral while simultaneously exhibiting altered domain organisation and condensation. Changes in mean radial position further supported the redistribution of euchromatin toward the nuclear periphery, particularly in stem and root under anaesthesia and combined treatments (**Figure 6f**). ^4,29,38^

**Figure 6.**
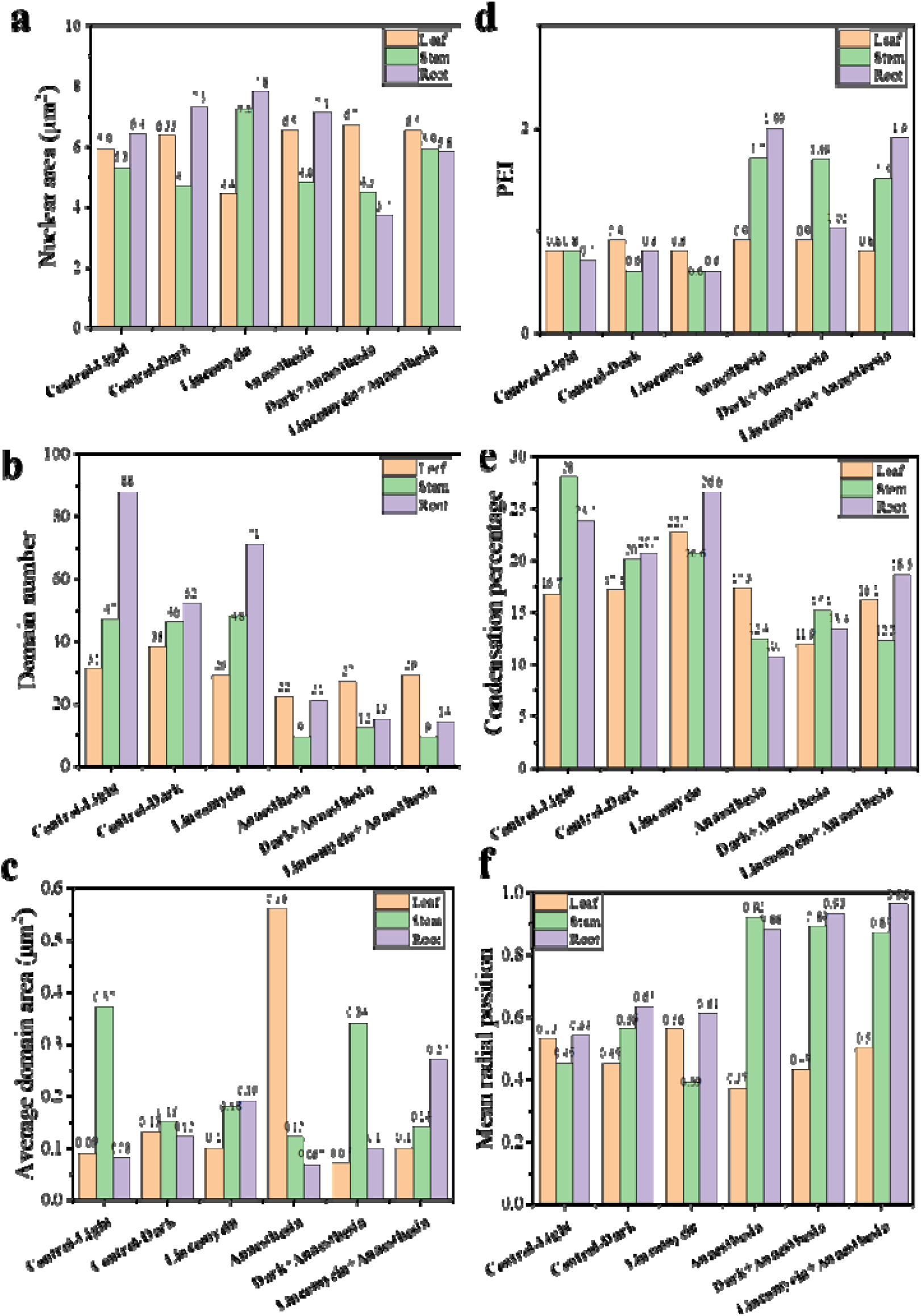
Quantitative characterization of euchromatin organization under the experimental conditions. **(a)** Nuclear area representing overall nuclear size. **(b)** Peripheral Enrichment Index (PEI), quantifying the relative enrichment of chromatin in the peripheral versus inner nuclear regions. **(c)** Number of chromatin domains representing the number of discrete chromatin domains detected within the nucleus. **(d)** Mean domain area representing the average area of individual chromatin domains. **(e)** Condensation percentage representing the proportion of nuclear area occupied by the analyzed chromatin domains. **(f)** Domain density representing the number of chromatin domains normalized to nuclear area.

The contrasting tissue responses suggest that anaesthesia induces a tissue-specific chromatin response rather than a uniform nuclear effect. The peripheral organisation observed in stem and root, supported by 3D Z-stack analysis, contrasts with the condensed euchromatin pattern in leaf and may reflect differences in tissue architecture and nuclear organisation. Leaf tissues contain loosely connected mesophyll cells separated by air spaces, while guard cells are structurally more isolated, potentially limiting long-range synchronisation of nuclear organisation. The occurrence of peripheral organisation in chloroplast-free root further suggests that anaesthesia-associated euchromatin redistribution can occur independently of chloroplast-derived signalling. In contrast, the distinct leaf response to lincomycin indicates that chloroplast functional status also contributes to euchromatin organisation. Thus, chloroplast-dependent and chloroplast-independent influences appear to converge on tissue-specific nuclear euchromatin architecture.

### 2.4 Heterochromatin Undergoes Tissue-Specific Spatial Inversion

Heterochromatin organisation, assessed by H3K9me3 immunofluorescence and SRRF imaging (**Figure 7**), revealed the most informative patterns in the dataset.^39^ The three tissues displayed distinct baseline heterochromatin architectures, and identical perturbations produced spatially opposite outcomes in different tissues, constituting a spatial inversion phenomenon. Quantitative parameters including nuclear area, domain number, mean domain area, Peripheral Enrichment Index (PEI), condensation index, and mean radial position are presented in **Figure 8**.

**Figure 7.**
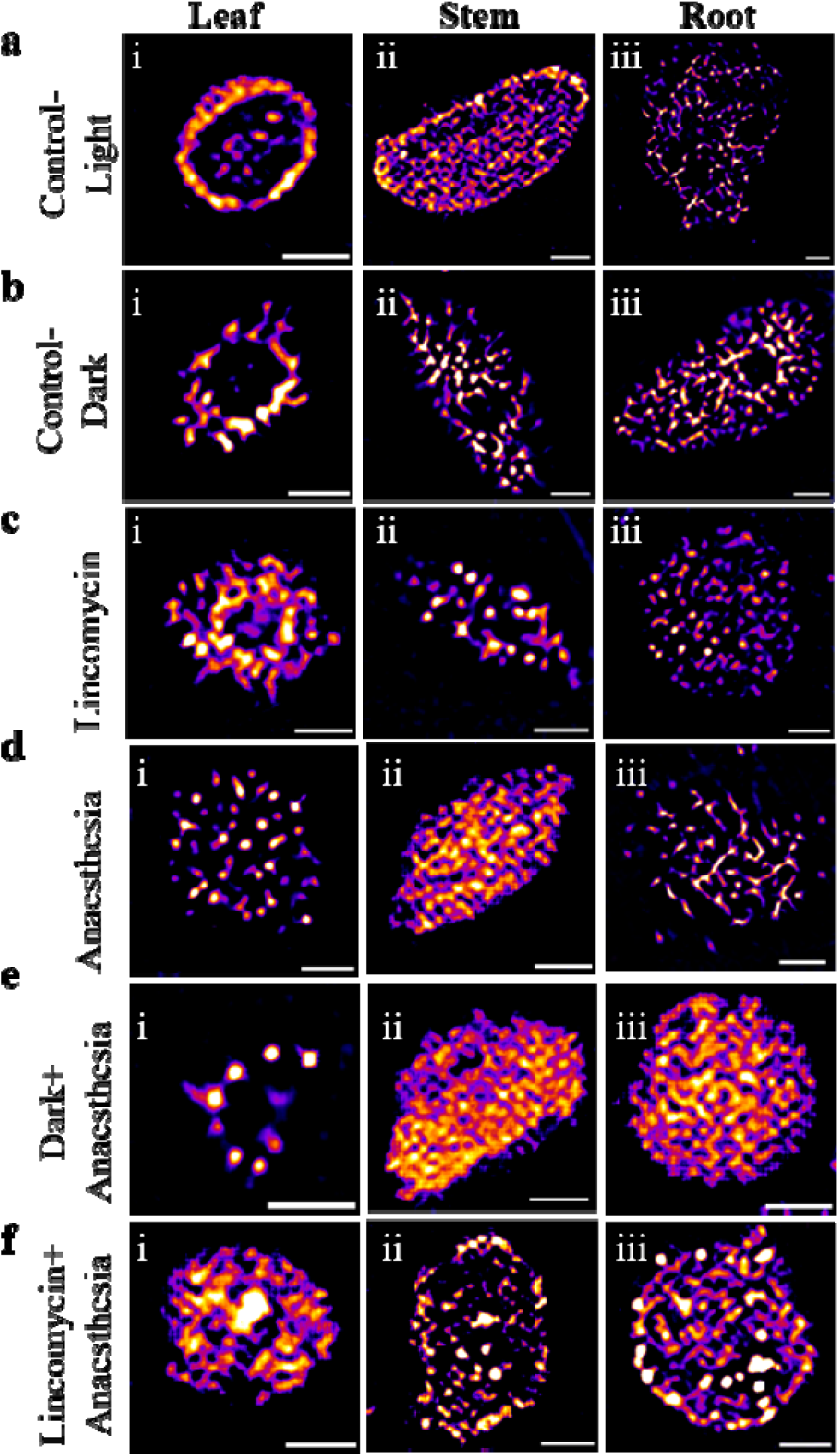
Representative SRRF images showing the spatial organisation of H3K9me3-marked heterochromatin across experimental conditions and tissues in *Solanum lycopersicum* seedlings. **(a-f)** correspond to **(a)** Control-Light, **(b)** Control-Dark, **(c)** Lincomycin, **(d)** Anaesthesia, **(e)** Dark + Anaesthesia, and **(f)** Lincomycin + Anaesthesia, respectively. Within each condition, **(i)** leaf, **(ii)** stem, and **(iii)** root nuclei are shown Control nuclei showed predominantly peripheral heterochromatin in leaf and a more dispersed organisation in stem and root. Lincomycin induced inward condensation in leaf but peripheral condensation in stem, with minimal change in root. Anaesthesia disrupted the leaf peripheral pattern and produced tissue-dependent condensation, while combined treatments produced central condensation in leaf and predominantly peripheral condensation in stem and root. Overall, the images demonstrate pronounced tissue- and condition-dependent heterochromatin reorganisation. Scale bars: 1000 nm.

**Figure 8.**
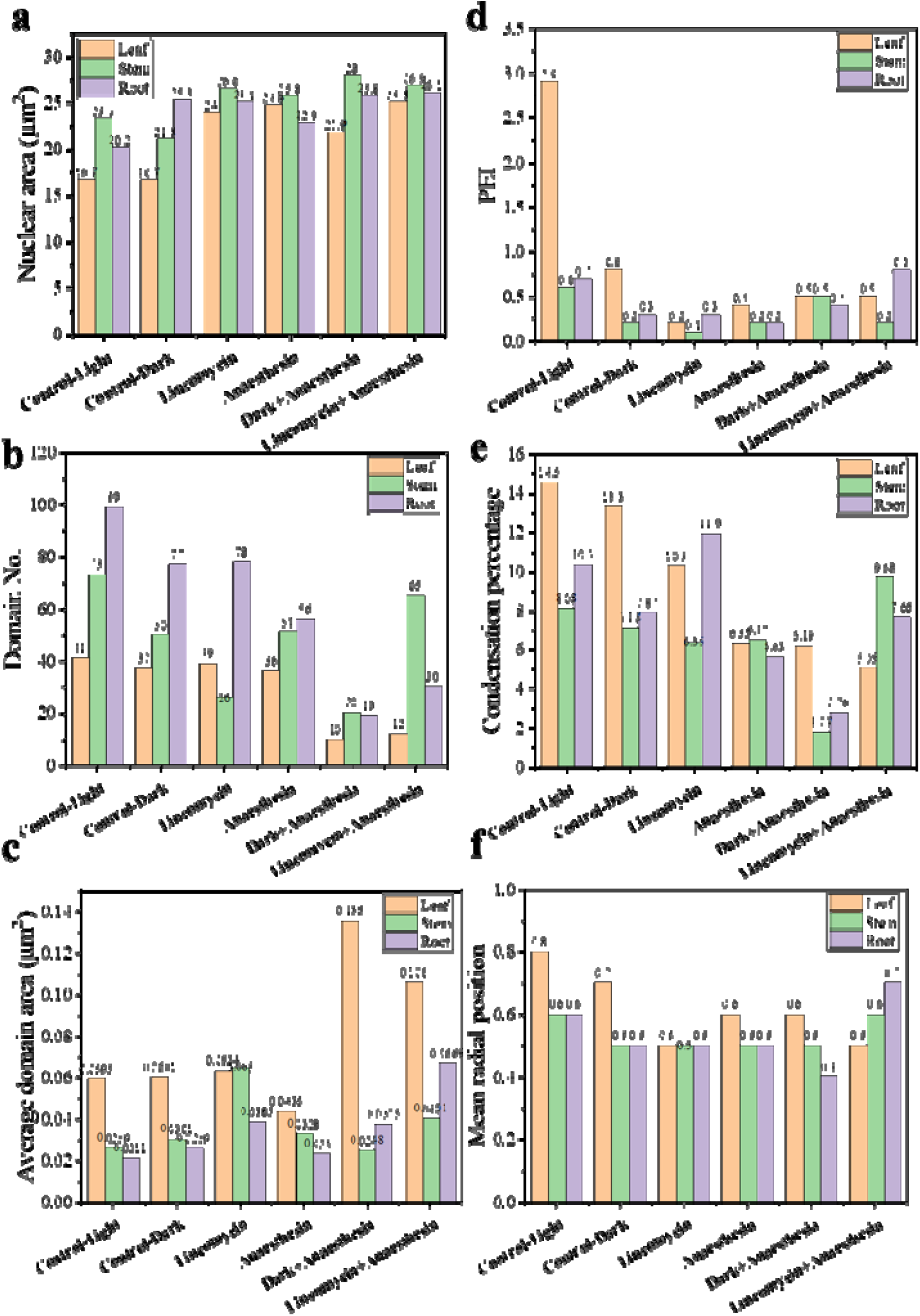
Quantitative characterization of heterochromatin organization under the experimental conditions. **(a)** Nuclear area representing overall nuclear size. **(b)** Peripheral Enrichment Index (PEI), indicating the relative enrichment of heterochromatin toward the nuclear periphery. **(c)** Number of heterochromatin domains representin the number of discrete heterochromatin foci within the nucleus. **(d)** Mean heterochromatin domain area representin the average size of individual heterochromatin domains. **(e)** Heterochromatin condensation percentage representin the proportion of nuclear area occupied by heterochromatin domains. **(f)** Heterochromatin domain density, representing the number of heterochromatin domains normalized to nuclear area.

Under Control-Light conditions, leaf nuclei displayed a characteristic peripheral heterochromatin ring, with H3K9me3-associated fluorescence concentrated at the nuclear boundary (**Figure 7**). Nuclear area was comparable across conditions within each tissue (**Figure 8a**), confirming that spatial chromatin differences were not attributable to nuclear size differences. The PEI was highest in Control-Light leaf (**Figure 8d**), domain number and mean domain area were consistent with well-organised chromocenters (**Figure 8b, c**), mean radial position confirmed domains were concentrated in the outermost nuclear shells (**Figure 8f**), and condensation index indicated a substantial proportion of nuclear area was occupied by heterochromatin domains (**Figure 8e**).^40,41^ This peripheral organisation reflects active maintenance of lamin-associated domains (LADs) at the inner nuclear membrane, driven by CRWN protein interactions and sustained by chloroplast-to-nucleus retrograde signals that reinforce the epigenetic state of heterochromatic loci.^5,42^ In contrast, Control-Light stem and root nuclei showed broadly distributed heterochromatin without a defined peripheral ring (**Figure 7**), with lower PEI values and more central mean radial positions (**Figure 8d, f**). Leaf cells maintain a high density of functional chloroplasts in close proximity to the nuclear envelope, providing a sustained localised source of retrograde signals that continuously reinforce peripheral heterochromatin anchoring. Stem and root cells receive this retrograde input at lower magnitude or not at all, defaulting to a distributed heterochromatin pattern. This gradient of peripheral organisation from leaf to stem to root directly parallels the gradient of chloroplast abundance and retrograde signalling capacity.^43,44^

Control-Dark conditions largely preserved the tissue-specific baseline organisation across all quantitative parameters (**Figure 8a-f**). This demonstrates that the peripheral heterochromatin ring in leaf nuclei is not maintained solely by light but reflects a developmental commitment established during chloroplast biogenesis. The etioplasts present in dark-grown tissue may provide residual retrograde signals possibly through labile tetrapyrrole intermediates or basal GUN1 activity sufficient to partially sustain peripheral chromatin architecture in the short term.^33,41,43^ Simple light deprivation is therefore insufficient to destabilise an established heterochromatin topology. A more fundamental disruption of plastid translational competence or retrograde signal generation is required.

Lincomycin treatment produced the first and most revealing demonstration of spatial inversion. In leaf nuclei, PEI and mean radial position both declined markedly (**Figure 8d, f**), confirming inward redistribution of heterochromatin (**Figure 7**). Condensation index declined while domain number showed modest change, indicating that interior-shifted domains were dispersed rather than compacted (**Figure 8b, e**). Without retrograde reinforcement, lamin-heterochromatin anchoring mediated by CRWN proteins weakens and H3K9me3-marked domains redistribute toward the nuclear interior.^5,12,44^. Crucially, this occurs while chloroplasts remain physically present, confirming that the functional output not the physical presence of the chloroplast determines nuclear heterochromatin topology. In stem nuclei, the same lincomycin treatment produced the opposite outcome: PEI and mean radial position increased (**Figure 8d, f**), domain number decreased while mean domain area increased (**Figure 8b, c**), indicating peripheral coalescence of heterochromatin domains (**Figure 7**). When retrograde signals from the limited stem chloroplasts are eliminated, residual heterochromatin associates with the nuclear membrane through default lamin-CRWN interactions independent of retrograde signalling. Stem therefore represents a system in which retrograde signals suppress peripheral heterochromatin whereas in leaf they promote it, a fundamentally opposite relationship producing spatial inversion under identical treatment. Root nuclei showed minimal change across all parameters under lincomycin alone (**Figure 8a-f**), as expected in the complete absence of chloroplasts. ^12,41,45^

Anaesthesia produced a distinct heterochromatin phenotype differing from both Control-Dark and lincomycin patterns. In leaf nuclei, PEI declined, domain number increased, mean domain area decreased, and mean radial position shifted to an intermediate value (**Figure 8b-d, f**), indicating broad dispersal throughout the nuclear volume rather than directional condensation (**Figure 7**). In stem nuclei, the opposite pattern emerged: reduced domain number, increased mean domain area, lower mean radial position, and reduced PEI (**Figure 8b-d, f**), indicating central condensation (**Figure 7**). Root nuclei showed only mild changes across all parameters (**Figure 8a-f**). ^4,36,46,47^

Under Dark + Anaesthesia, leaf nuclei showed the lowest domain number and highest maximum domain area in the dataset (**Figure 8b, c**), indicating substantial chromocenter fusion, while stem and root showed interior compaction and markedly reduced condensation index (**Figure 8e**). Under Lincomycin + Anaesthesia, leaf heterochromatin condensed toward the interior with reduced PEI and radial position (**Figure 8d, f**), while stem and root showed strong peripheral organisation with elevated PEI, with the root value representing the second highest in the entire dataset despite roots lacking chloroplasts. Together, these results establish that heterochromatin undergoes highly tissue-specific and condition-dependent spatial reorganisation, and that the direction of heterochromatin relocalization is determined by the tissue-specific relationship between chloroplast retrograde signalling status and baseline nuclear chromatin architecture. ^42^

### 2.5 Chloroplast Retrograde Signalling and Nuclear Envelope Mechanics Jointly Determine Nuclear Chromatin Topology in a Tissue-Specific Manner

The spatial organisation of euchromatin and heterochromatin in *Solanum lycopersicum* nuclei is governed by two converging regulatory inputs: chloroplast retrograde signalling and a chloroplast-independent pathway sensitive to anaesthetic exposure, both acting in a tissue-specific manner.^48^ Active chloroplasts in leaf mesophyll cells generate a continuous flux of retrograde signal molecules like ROS, PAP, MEcPP, and tetrapyrrole intermediates that are delivered to the nucleus at high local concentration owing to close chloroplast-nuclear proximity. ^10,49,50^ These signals engage the GUN1 network, stabilise H3K9me3-marked heterochromatin at the nuclear periphery through CRWN protein-LAD interactions, and maintain H3K4me3-marked euchromatin in spatially confined transcriptionally active domains.^5,40^ When this retrograde signal is attenuated by darkness or eliminated by lincomycin, leaf heterochromatin detaches from the nuclear periphery and redistributes inward, while stem heterochromatin moves in the opposite direction toward the periphery because retrograde signals were suppressing rather than promoting peripheral organisation in stem.^4,46^ Root nuclei, lacking chloroplasts entirely, show minimal response to lincomycin, confirming that this axis is chloroplast-dependent.

Anaesthesia produces qualitatively distinct chromatin phenotypes that occur even in chloroplast-free root cells, establishing a plastid-independent pathway. Anaesthetic agents alter nuclear envelope membrane properties and the mechanical state of inner nuclear membrane proteins, disrupting the tension environment that maintains chromatin domain positioning at the lamina.^4,36^ This produces dispersal in leaf and central condensation in stem through NAD interactions, independently of any change in retrograde signalling. ^51^Under combined perturbations, both inputs are disrupted simultaneously, producing non-additive outcomes including chromocenter fusion in leaf and near-uniform interior compaction in stem and root. Nuclear chromatin topology therefore encodes the integrated state of organellar signalling and nuclear envelope physical properties, functioning as a spatial biomarker of cellular perturbation state in plants.^52^

## 3. Conclusion

This study demonstrates that nuclear chromatin organisation in *Solanum lycopersicum* is strongly influenced by tissue context and experimental perturbation. Confocal and SRRF imaging revealed distinct, tissue-dependent changes in euchromatin and heterochromatin following chloroplast perturbation and anaesthetic exposure. Lincomycin produced contrasting heterochromatin redistribution in leaf and stem, while root nuclei showed comparatively limited changes under lincomycin alone. In contrast, anaesthesia induced pronounced peripheral organisation of euchromatin, particularly in stem and root, and this phenotype persisted under combined anaesthesia with darkness or lincomycin. These findings indicate that anaesthesia-associated chromatin reorganisation is not solely dependent on normal chloroplast function and includes a chloroplast-independent component. The quantitative spatial framework provides a systematic approach for distinguishing tissue- and condition-specific chromatin phenotypes. Although the observed patterns are consistent with involvement of chloroplast-to-nucleus signalling, the underlying molecular mechanisms require further investigation. Overall, this study establishes nuclear chromatin topology as a sensitive spatial readout of plant responses to chloroplast perturbation and anaesthesia.

## Author Contributions

SC conceived and designed the experiments with inputs from CKN and LB, optimized experimental protocols, performed all plant-related experiments, imaging, and data analysis, and also wrote the manuscript. SCH helped in plant related experiments and imaging. CKN and LB supervised the overall project, offered continuous guidance throughout the research, and contributed to the conceptual development and editing of the manuscript.

## Data Availability Statement

The data that support the findings of this study are available in the supplementary material of this article.

## Acknowledgements

The authors acknowledge the facilities and technical assistance of the Indian Institute of Technology Mandi and the support of the IKSMHA Centre. We thank AMRC for the confocal and SRRF imaging facility for instrument access.

