## Supplementary Information for "Euchromatin Peripheral Organization Follows Anterograde Signalling Under Anaesthetic Stress"

### Materials and Methods

#### Plant Material and Experimental Conditions

Fourteen-day-old plants were used for all experiments. Plants were maintained under controlled growth conditions on ½× Murashige and Skoog (MS) medium supplemented with 1% sucrose at 22-24°C. Six experimental conditions were established: Control-Light, Control-Dark, Lincomycin, Anaesthesia, Dark + Anaesthesia, and Lincomycin + Anaesthesia. Control-Light plants were maintained under the standard light condition, whereas Control-Dark plants were maintained in darkness. For chloroplast perturbation, plants were treated with lincomycin under the corresponding growth conditions. Lincomycin treatment was used to interfere with plastid translation and chloroplast development. For anaesthesia experiments, plants were exposed to 1% (w/v) lidocaine for 1 h. Anaesthesia was applied independently to generate the Anaesthesia condition and in combination with the preceding dark or lincomycin treatment to generate the Dark + Anaesthesia and Lincomycin + Anaesthesia conditions.

At the completion of the experimental treatment, plants were collected for phenotypic characterization, pigment analysis, chloroplast imaging, nuclear imaging, and chromatin immunofluorescence analysis. The experimental design was intended to distinguish effects associated with light availability and chloroplast perturbation from those associated with anaesthesia and to determine whether these perturbations produced tissue-specific changes in nuclear organization.

#### Phenotypic Characterization

Whole-plant morphology was documented for each experimental condition. Plant growth was quantified using leaf length, stem length, root length, and leaf number as phenotypic parameters. Leaf, stem, and root lengths were measured in centimetres, whereas leaf number was recorded as the total number of leaves per plant.

For each experimental condition, 20 individual plants (n = 20) were analyzed. Representative images were acquired to document overall morphological differences, and quantitative measurements were used to assess condition-dependent changes in shoot and root development.

**Pigment Content Analysis by UV-Visible Spectrophotometry**

Photosynthetic pigment content was quantified to assess treatment-associated changes in chloroplast function. Chlorophyll a, chlorophyll b, and carotenoid contents were determined from fresh leaf tissue using acetone extraction followed by UV-visible spectrophotometry.

For each sample, exactly 52.5 mg of fresh leaf tissue was weighed using an analytical balance and transferred to a chilled mortar. The tissue was thoroughly homogenized using a pestle in 2 mL of 80% acetone until a uniform green slurry was obtained. The homogenate was transferred to a 15-mL sterile centrifuge tube and centrifuged at 8,000-10,000 rpm for 10 min at 4°C using an Eppendorf 5804R tabletop centrifuge.

Following centrifugation, the clear supernatant was carefully transferred to a fresh tube without disturbing the pellet. Absorbance was measured using a Thermo Scientific Genesys 10S UV-Vis spectrophotometer, with 80% acetone used as the blank. Spectral measurements were acquired over 400-700 nm. Absorbance values at 663, 645, and 470 nm were recorded for chlorophyll a, chlorophyll b, and carotenoids, respectively. Pigment concentrations were calculated using standard extinction coefficients.

**Tissue Collection and Fluorescence Preparation**

Leaf, stem, and root tissues were collected for cellular and nuclear analyses. The three tissues were selected to allow comparison of chloroplast-containing and predominantly non-photosynthetic tissues and to determine whether nuclear responses to the experimental conditions were tissue dependent.

For nuclear DNA visualization, DAPI-containing mounting medium was used where applicable. For chromatin-state analysis, tissue samples were subjected to immunofluorescence labeling of histone modifications associated with euchromatin and heterochromatin.

**Chromatin Immunofluorescence Staining**

Histone modification-based chromatin states were assessed by immunofluorescence staining. H3K4me3 was used as a marker of actively transcribed euchromatin, whereas H3K9me3 was used as a marker of transcriptionally silent heterochromatin. Tissue sections were incubated overnight at 4°C with primary antibodies against H3K4me3 and H3K9me3. Both primary antibodies were used at a 1:100 dilution. The H3K4me3 antibody was Abclonal A4097, rabbit monoclonal, and the H3K9me3 antibody was Abclonal A4098, rabbit monoclonal. Following primary antibody incubation, samples were washed three times for 5 min each with PBS containing 0.1% Tween-20 (PBS-T) to remove unbound antibody. A Cy3-conjugated goat anti-rabbit secondary antibody (Abclonal A0520, or equivalent) was applied at a 1:600 dilution for 1 h at room temperature in darkness. After secondary antibody incubation, samples were washed three additional times for 5 min with PBS-T and subsequently rinsed three times with

PBS. Coverslips were mounted using either glycerol:PBS (1:1, v/v) mounting medium containing DAPI. Coverslips were sealed with nail polish to prevent dehydration during imaging.

Positive and negative controls were included in each staining batch. Negative controls consisted of tissue processed without primary antibody and tissue exposed to secondary antibody alone. These controls were used to assess nonspecific fluorescence and verify the specificity of antibody-associated fluorescence.

**Confocal Laser Scanning Microscopy**

Confocal imaging was performed using a Nikon Eclipse Ti inverted microscope. Images were acquired with Nikon NIS-Element software. Cell samples were excited with a 405, 488, 561, and 639 nm laser, and emissions were collected using appropriate filter sets. Most images were acquired using a 60×, 1.40 NA oil-immersion Nikon Plan Apo objective with Immersol 518F immersion oil. Three-dimensional Z-stack images were acquired using 1-µm optical slice intervals. The Z-stack acquisition range was approximately 2350-2400 nm to accommodate imaging of the complete nuclear and organellar volume. Images were acquired at 512 × 512-pixel resolution with 8-bit depth at room temperature (20-22°C). The confocal pinhole was set to 1.0 Airy unit. Image averaging was set to two frames per slice to reduce image noise while preserving spatial information.

**Super-Resolution Radial Fluctuation Imaging**

Super-resolution radial fluctuation (SRRF) imaging was used to visualize euchromatin and heterochromatin at enhanced spatial resolution. SRRF imaging was performed using a Nikon Ti Eclipse microscope equipped with a 100× Plan Apo λ oil-immersion objective (NA 1.45) and an additional 1.5× magnification lens. Images were acquired using an Andor iXon Ultra 897U EMCCD camera with a 1024 × 1024-pixel sensor. The camera was operated at an exposure time of 50 ms, corresponding to approximately 20 frames per second, with a 17-MHz readout rate and 16-bit acquisition depth. The electron multiplication gain was set to 3 on the preamplifier.

The Nikon Perfect Focus System was used during acquisition to maintain focus and minimize focal drift during the extended time-series acquisition required for SRRF reconstruction. For SRRF reconstruction, movies comprising 3,000 sequential frames were acquired. An additional 1,000 frames were acquired in temporal mode from the same optical field. Image sequences were saved in FITS format for subsequent processing. H3K4me3- and H3K9me3-associated fluorescence was acquired using the appropriate excitation and emission filter combinations for the fluorophore used.

**SRRF Image Reconstruction and Processing**

SRRF reconstruction was performed using the NanoJ-SRRF plugin in ImageJ/Fiji. Time-series datasets comprising 3,000-5,000 frames were acquired for each sample under equivalent imaging conditions.

The SRRF reconstruction parameters were established through preliminary optimization experiments. The parameters used for reconstruction were a ring radius of 0.5 pixels, radiality magnification of 5, and six axes in the ring. These parameters were selected to provide improved structural definition while minimizing reconstruction artifacts.

During reconstruction, each original pixel was subdivided into a 5 × 5-pixel grid, resulting in a fivefold increase in spatial sampling in the x and y dimensions. Background artifacts were removed using identical processing parameters across samples within a condition. Reconstructed images were subsequently scaled and merged in ImageJ/Fiji to generate the final super-resolved images used for visualization and quantitative analysis.

**Image Processing and Nuclear Segmentation**

Image processing and quantitative analysis were performed using ImageJ/Fiji version 1.53k (National Institutes of Health). For visualization of three-dimensional confocal datasets, maximum-intensity projections were generated using the Z Project function with the maximum-intensity projection option. This approach generated two-dimensional representations containing the maximum fluorescence intensity recorded at each xy-coordinate throughout the acquired Z-stack.

**Chloroplast Visualization and Quantification**

Chloroplasts were visualized using their intrinsic chlorophyll autofluorescence. Chloroplast fluorescence was acquired using the 488-nm excitation channel during confocal imaging. Leaf and stem tissues were analyzed to characterize treatment-associated changes in chloroplast organization. Chloroplast structures were identified from their fluorescence signal and quantified with respect to number and area. These parameters were used to assess changes in chloroplast abundance and morphology across the six experimental conditions.

Chloroplast fluorescence images were also examined together with DAPI fluorescence to assess the spatial association between chloroplasts and nuclei. For three-dimensional imaging, confocal Z-stacks were acquired through the tissue to capture the spatial relationship between chloroplasts and nuclei.

**Chloroplast-Nucleus Association Analysis**

The spatial relationship between chloroplasts and nuclei was evaluated in leaf and stem tissues using merged fluorescence images containing chloroplast autofluorescence and nuclear DNA fluorescence. Individual optical sections and maximum-intensity projections were examined to characterize the apparent spatial relationship between chloroplasts and nuclei. Z-stack datasets were used to distinguish spatial association observed in individual optical sections from the three-dimensional organization captured across the nuclear volume.

The analysis focused on condition-dependent changes in chloroplast number, size, distribution, and their spatial association with the nucleus.

**Quantitative Analysis of Nuclear Chromatin Organization**

Chromatin organization was quantified using spatial, morphological, and intensity-based measurements. The analysis was designed to characterize not only the amount of chromatin present within the nucleus but also its distribution, domain organization, and spatial relationship to the nuclear boundary. The principal measurements included nuclear area, Peripheral Enrichment Index, chromatin-domain number, mean domain area, condensation percentage, mean domain circularity, mean radial position, inter-domain nearest-neighbor distance, and euchromatin intensity distribution.

**Nuclear Area**

The total area enclosed by the segmented nuclear boundary was measured in **µm²**. Nuclear area was used as an indicator of nuclear size and as the normalization parameter for measurements involving chromatin-domain abundance and density.

**Peripheral Enrichment Index**

The **Peripheral Enrichment Index (PEI)** was calculated to quantify the relative enrichment of chromatin fluorescence at the nuclear periphery:

where **R** represents the nuclear radius.

$$PEI = \frac{\text{Mean intensity}_{0.8R-1.0R}}{\text{Mean intensity}_{0-0.6R}}$$

The numerator represents the mean fluorescence intensity in the outer 20% of the nuclear radius, corresponding to **0.8R-1.0R**, whereas the denominator represents the mean fluorescence intensity in the inner 60% of the nuclear radius, corresponding to **0-0.6R**.

A PEI greater than 1 indicates relative enrichment of the analyzed chromatin signal toward the nuclear periphery, whereas a PEI below 1 indicates relatively greater signal toward the nuclear interior.

**Chromatin-Domain Number and Mean Domain Area**

The number of discrete chromatin domains was determined using connected-component analysis of the segmented fluorescence signal. The mean domain area (µm²) was calculated as the mean area of all detected chromatin domains within each nucleus.

These parameters were interpreted jointly because the number and size of domains provide complementary information about chromatin organization. A large number of small domains indicates a more fragmented organization, whereas fewer larger domains are consistent with greater domain coalescence or fusion.

#### Chromatin Condensation Percentage

The proportion of the nuclear area occupied by segmented chromatin domains was calculated as:

$$\text{Condensation } (\%) = \frac{\text{Total area of chromatin domains}}{\text{Total nuclear area}} \times 100$$

This parameter represents the relative nuclear area occupied by the analyzed chromatin state.

#### Mean Radial Position

The spatial position of each chromatin-domain centroid was normalized to the nuclear radius. A normalized radial position of **0** corresponds to the nuclear center, whereas **1** corresponds to the nuclear periphery.

The mean radial position was calculated from the normalized centroid positions of all detected domains within an individual nucleus. This parameter was used to determine whether chromatin domains were preferentially localized toward the nuclear interior or periphery.

#### Reproducibility and Sample Size

Phenotypic measurements were obtained from 20 individual plants per experimental condition (n = 20). Microscopy-based quantitative analyses were performed using 10 nuclei per experimental condition (n = 10). Image acquisition and processing parameters were kept consistent across experimental conditions. The same segmentation criteria, radial analysis procedure, and quantitative definitions were applied across the dataset to permit direct comparison of nuclear, euchromatin, heterochromatin, and chloroplast organization.

#### Statistical Analysis

Quantitative data were analyzed by comparing the experimental conditions for each phenotypic, chloroplast, nuclear, and chromatin parameter. Statistical analyses were performed using the statistical approach appropriate for the distribution and experimental structure of each dataset. Multiple comparisons were corrected where applicable.

The statistical significance threshold, specific statistical tests, post-hoc procedures, and multiple-comparison correction should be reported exactly according to the statistical analyses performed for the final dataset. Individual observations were retained in the graphical representation wherever applicable to show the distribution of the experimental measurements.

### Supplementary Figures

**Figure S1.** Plant growth and photosynthetic pigment changes under the experimental conditions.

**Figure S2.** Three-dimensional confocal imaging of chloroplast-nucleus organization in leaf tissue.

**Figure S3.** Three-dimensional confocal imaging of chloroplast-nucleus organization in stem tissue.

**Figures S4-S9.** Confocal z-stack imaging of chloroplast-nucleus organization in leaf tissue under Control-Light, Control-Dark, Lincomycin, Anaesthesia, Dark + Anaesthesia, and Lincomycin + Anaesthesia conditions, respectively.

**Figures S10-S15.** Confocal z-stack imaging of chloroplast-nucleus organization in stem tissue under Control-Light, Control-Dark, Lincomycin, Anaesthesia, Dark + Anaesthesia, and Lincomycin + Anaesthesia conditions, respectively.

**Figures S16-S21.** Confocal z-stack imaging of nuclear DNA organization in root tissue under Control-Light, Control-Dark, Lincomycin, Anaesthesia, Dark + Anaesthesia, and Lincomycin + Anaesthesia conditions, respectively.

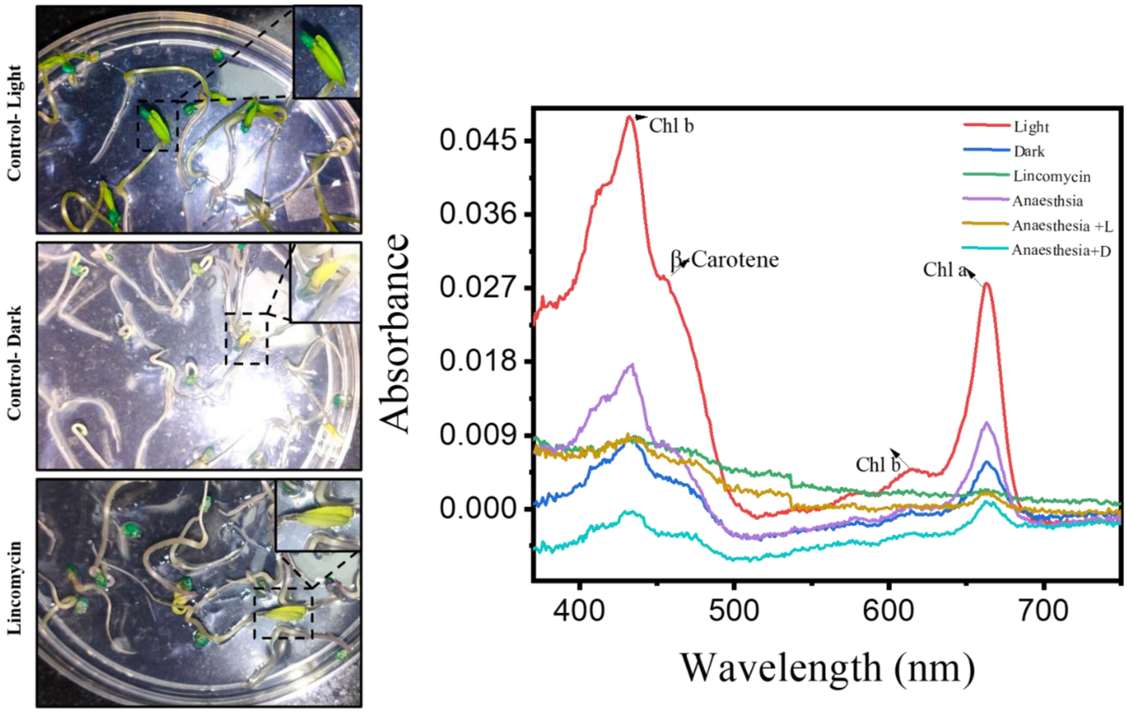


**Figure S1. Plant growth and pigment changes under different stress conditions.** Representative photographs of seedlings grown under different experimental conditions, with boxed regions shown as magnified views. Lincomycin-treated and dark-grown seedlings exhibit pronounced pigment disturbances, characterized by reduced greening and yellowing of the leaves, indicating impaired chloroplast development and pigment accumulation. The corresponding UV-visible absorption spectra show a reduction in absorbance under different stress conditions compared with the control, consistent with decreased photosynthetic pigment content. Different coloured traces represent the respective growth/stress conditions.

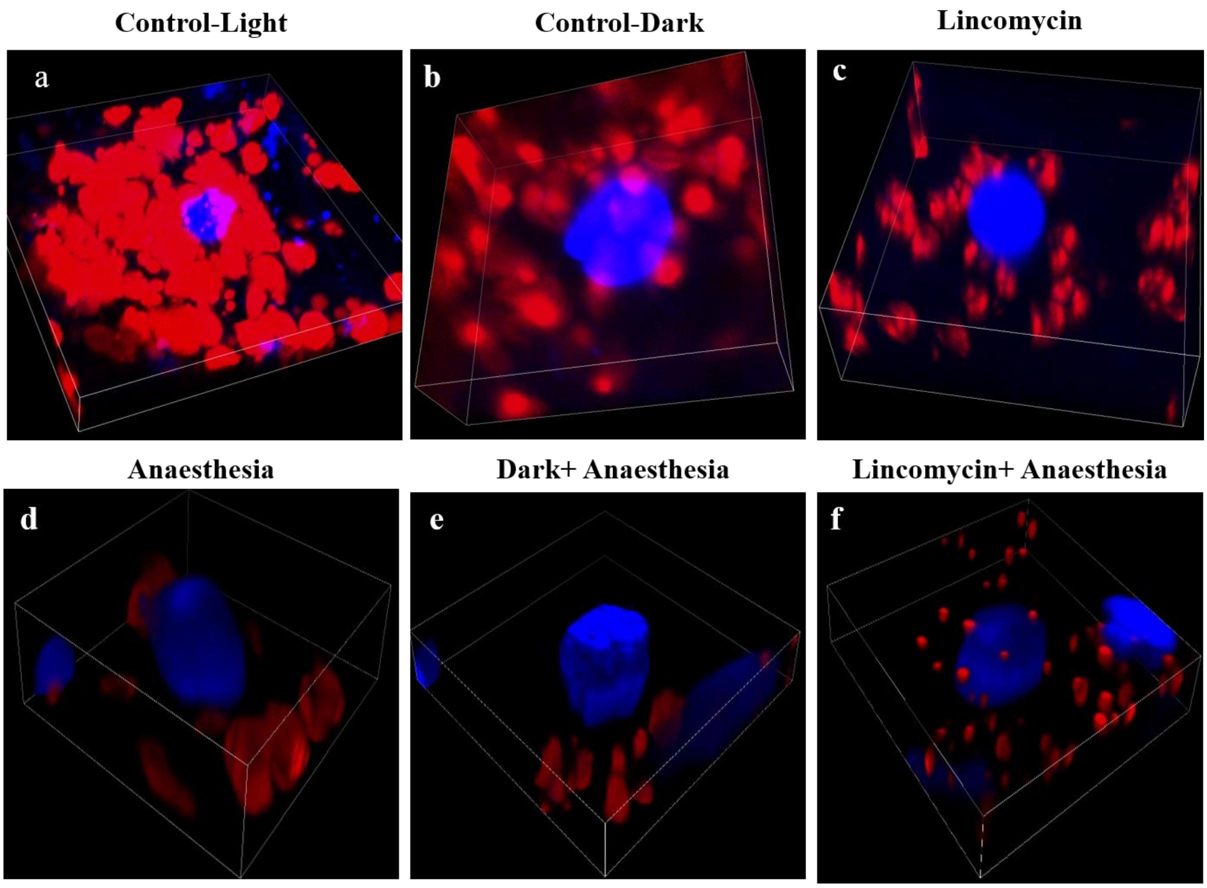


**Figure S2. Three-dimensional confocal fluorescence imaging of chloroplast and nuclear organization of leaf under different growth conditions.** Representative 3D reconstructed confocal images (a-f) showing the spatial distribution and organization of chloroplast-associated red autofluorescence and nuclear blue fluorescence under different experimental conditions. The images reveal condition-dependent changes in chloroplast abundance, distribution, and spatial association with the nucleus. Red fluorescence represents chloroplast/pigment autofluorescence, whereas blue fluorescence represents nuclear staining. Differences in fluorescence intensity and chloroplast organization across the conditions indicate altered chloroplast development and cellular organization under stress or modified growth conditions.

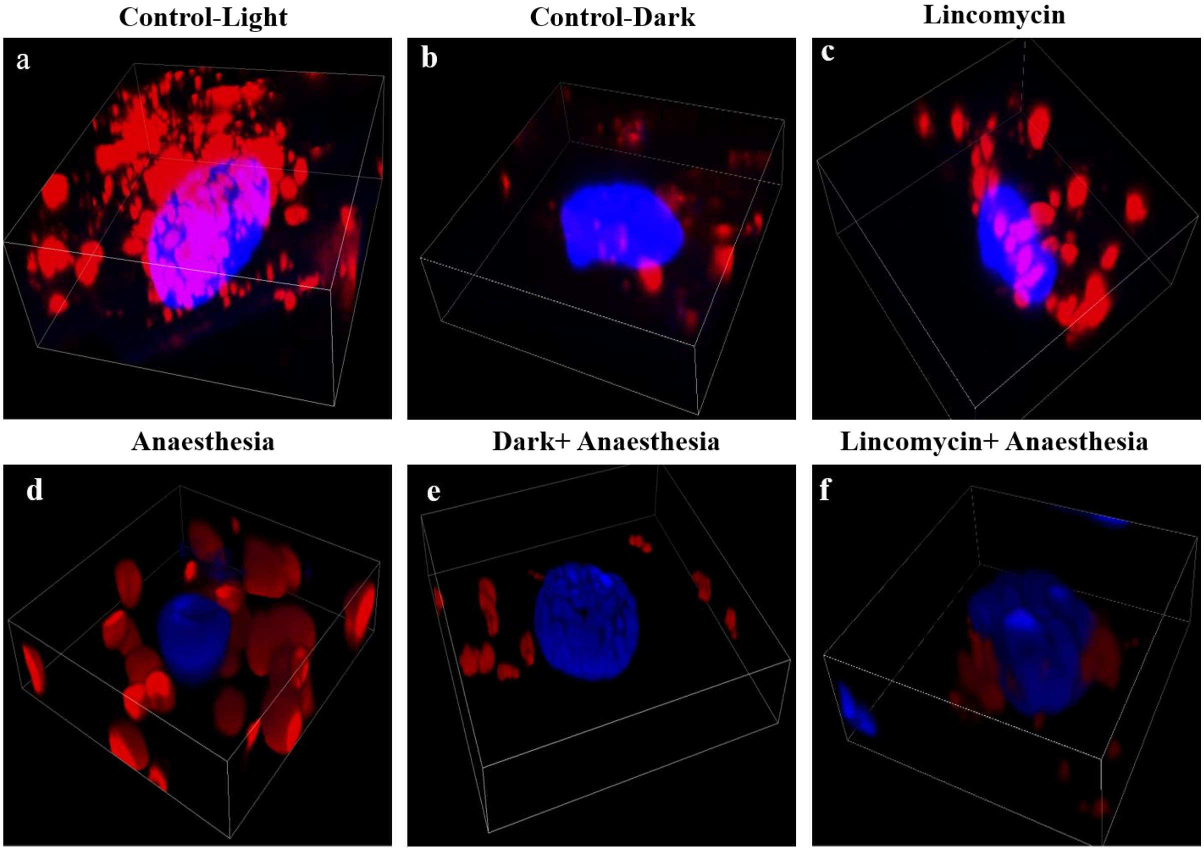


**Figure S3. Three-dimensional confocal reconstruction showing chloroplast-nucleus organization of stem under different experimental conditions.** Representative 3D confocal images (a-f) illustrating the spatial distribution of chloroplasts (red autofluorescence) and nuclei (blue fluorescence) under different growth or treatment conditions. The images demonstrate condition-dependent changes in chloroplast abundance, morphology, distribution, and their spatial relationship with the nucleus. Variations in red fluorescence intensity and chloroplast organization indicate altered pigment accumulation and chloroplast development, while changes in nuclear morphology and chloroplast-nucleus association reflect alterations in cellular organization under the respective conditions.

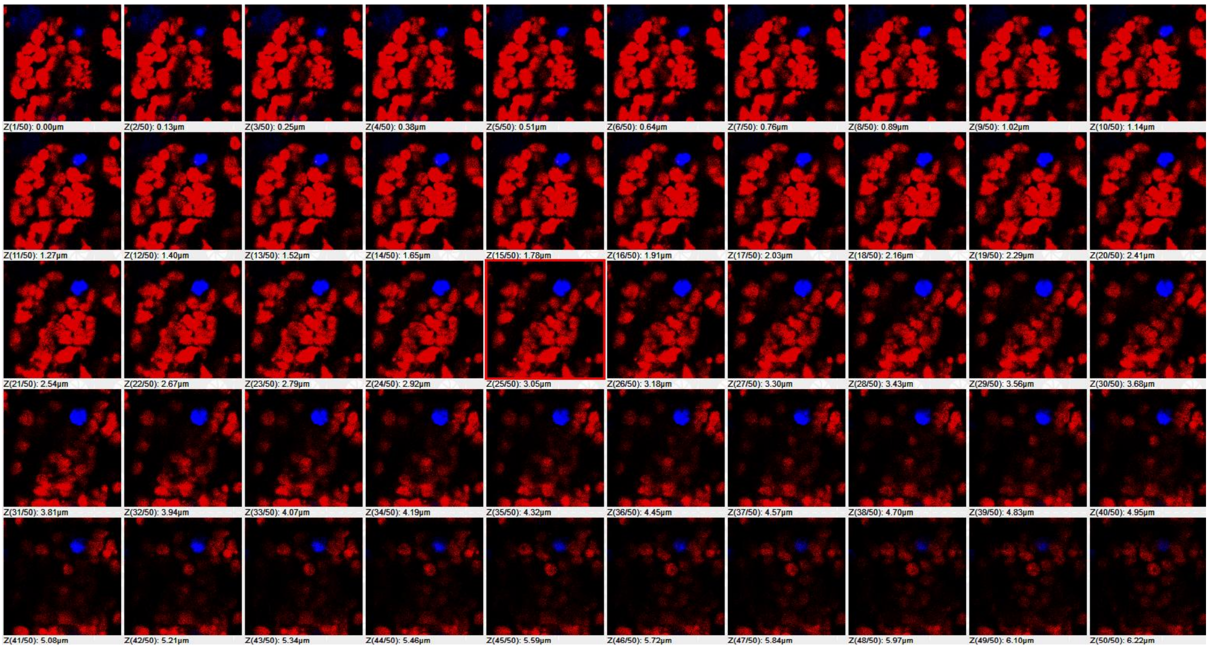


**Figure S4. Confocal z-stack analysis of chloroplast-nucleus organization in leaf tissue under control light conditions.** Representative sequential confocal optical sections (Z1-Z50) through a leaf tissue z-stack showing the spatial distribution of chloroplasts and nuclei under control light conditions. Red fluorescence represents chloroplast autofluorescence, while blue fluorescence represents nuclear staining. The individual optical sections are presented sequentially along the z-axis demonstrating the three-dimensional distribution and spatial organization of chloroplasts relative to the nucleus within the leaf tissue.

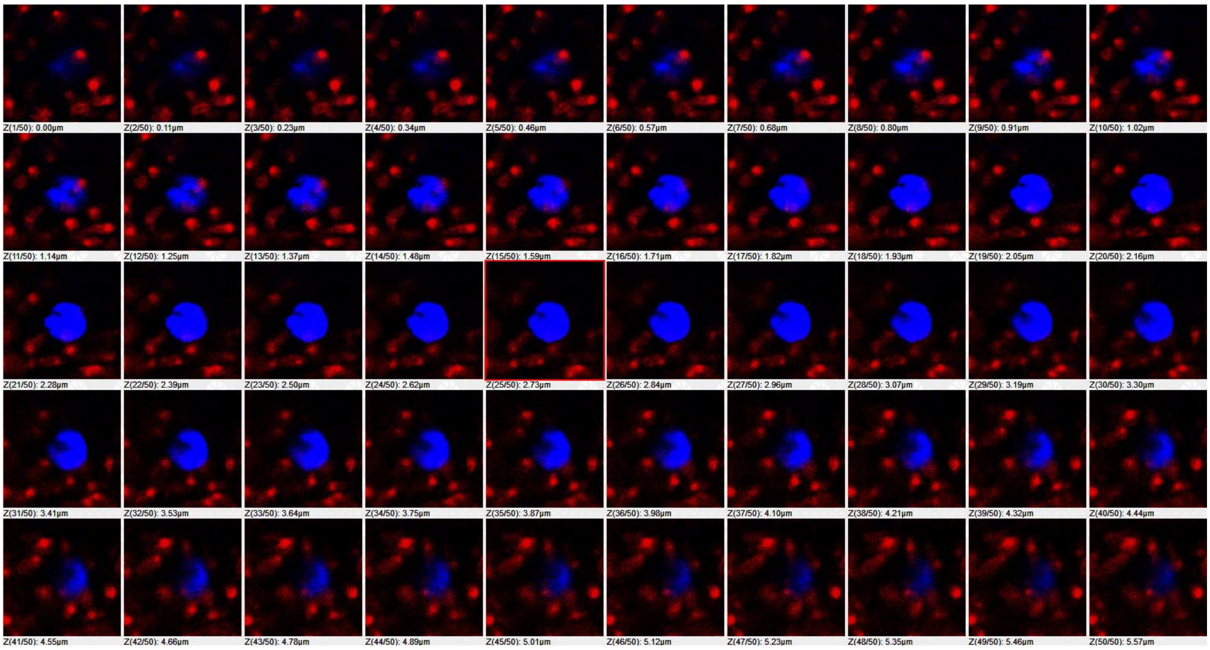


**Figure S5. Confocal z-stack imaging of chloroplast-nucleus organization in dark-grown leaf tissue.** Representative sequential confocal optical sections (Z1-Z50) acquired through dark-grown leaf tissue. Red fluorescence represents chloroplast autofluorescence, while blue fluorescence represents nuclear staining. The sequential z-sections demonstrate the spatial distribution of chloroplasts surrounding the nucleus and the altered chloroplast fluorescence and organization associated with dark-grown conditions. Reduced chloroplast/pigment-associated fluorescence is consistent with impaired chloroplast development and pigment accumulation under darkness.

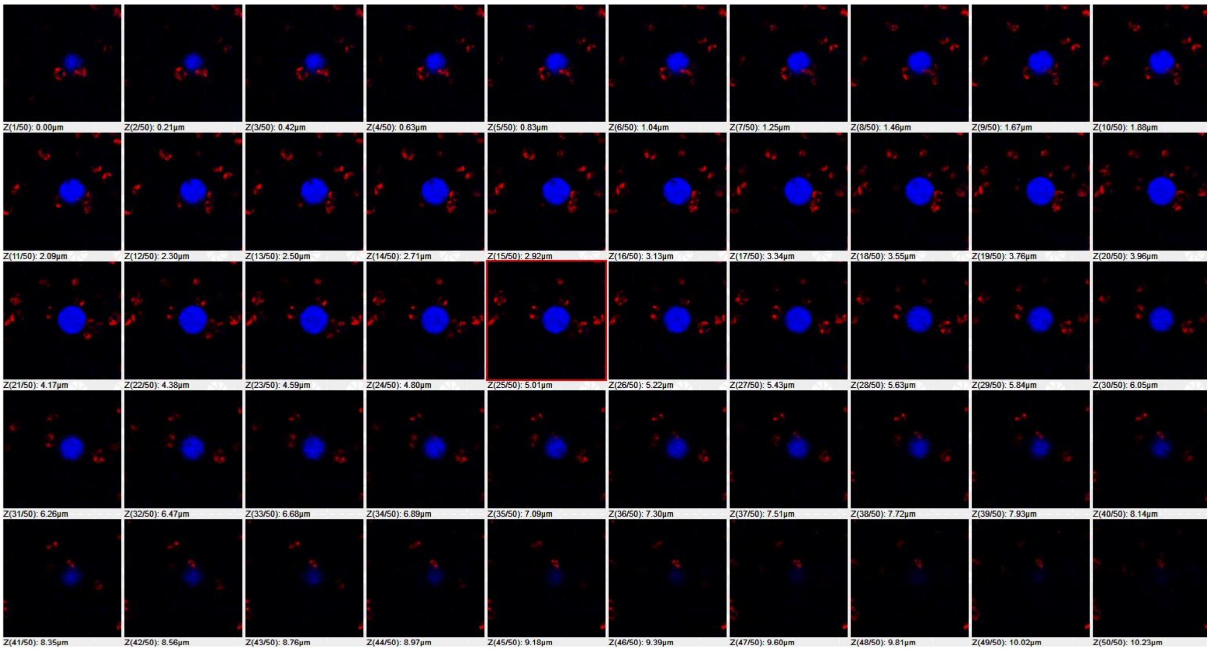


**Figure S6. Confocal z-stack imaging of chloroplast-nucleus organization in lincomycin-grown leaf tissue.** Representative sequential confocal optical sections (**Z1-Z50**) acquired through lincomycin-treated leaf tissue. Red fluorescence represents chloroplast autofluorescence, while blue fluorescence represents nuclear staining. The z-stack demonstrates the spatial distribution of chloroplasts surrounding the nucleus and the altered chloroplast-associated fluorescence following inhibition of chloroplast development by lincomycin.

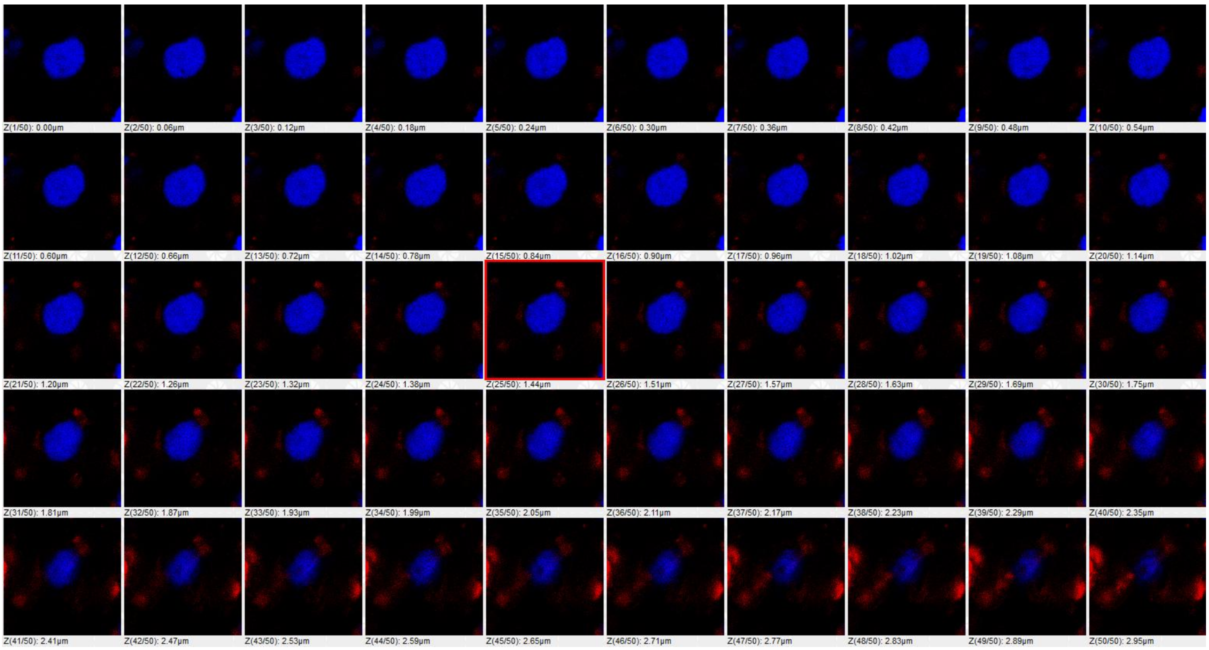


**Figure S7. Confocal z-stack imaging of chloroplast-nucleus organization in anaesthesia-treated leaf tissue.** Representative sequential confocal optical sections (**Z1-Z50**) acquired through anaesthesia-treated leaf tissue. Red fluorescence represents chloroplast autofluorescence, whereas blue fluorescence represents nuclear staining. The z-stack illustrates changes in chloroplast distribution and fluorescence intensity and their spatial relationship with the nucleus following anaesthesia treatment, indicating altered chloroplast organization and pigment-associated fluorescence.

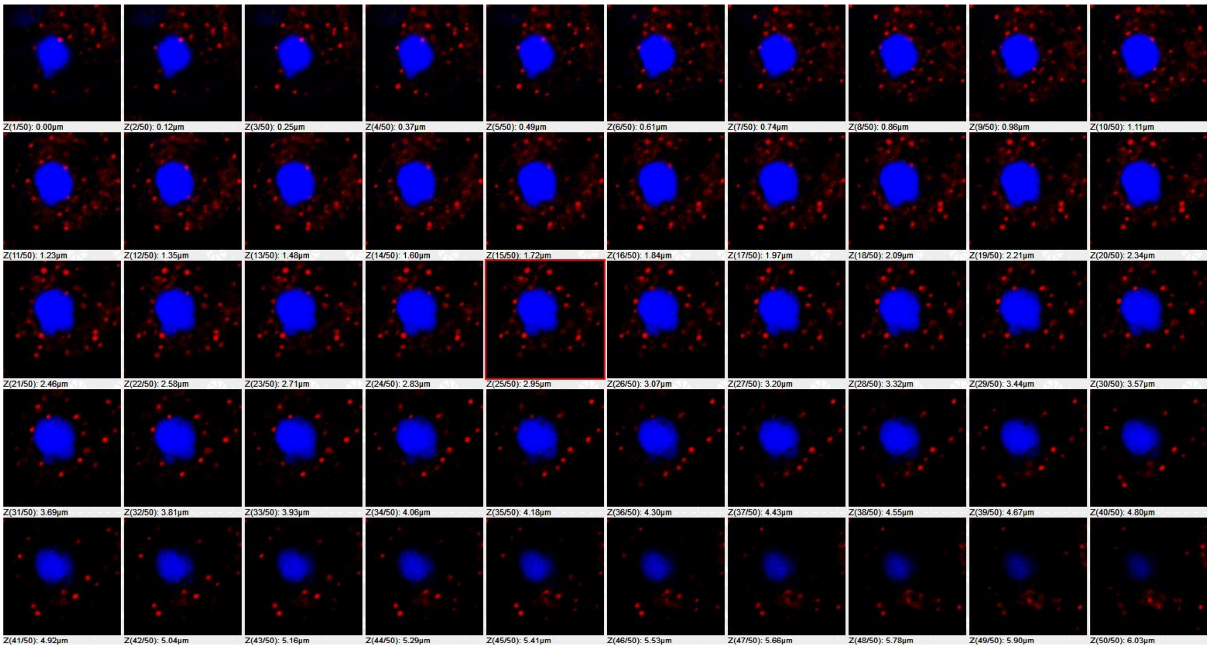


**Figure S8. Confocal z-stack imaging of chloroplast-nucleus organization in dark-grown, anaesthesia-treated leaf tissue.** Representative sequential confocal optical sections (**Z1-Z50**) showing chloroplast and nuclear organization in leaf tissue exposed to darkness and anaesthesia. Red fluorescence represents chloroplast autofluorescence, while blue fluorescence represents nuclear staining. The z-stack reveals reduced chloroplast-associated fluorescence and altered chloroplast distribution around the nucleus, reflecting the combined effects of dark growth and anaesthesia on chloroplast development and pigment accumulation.

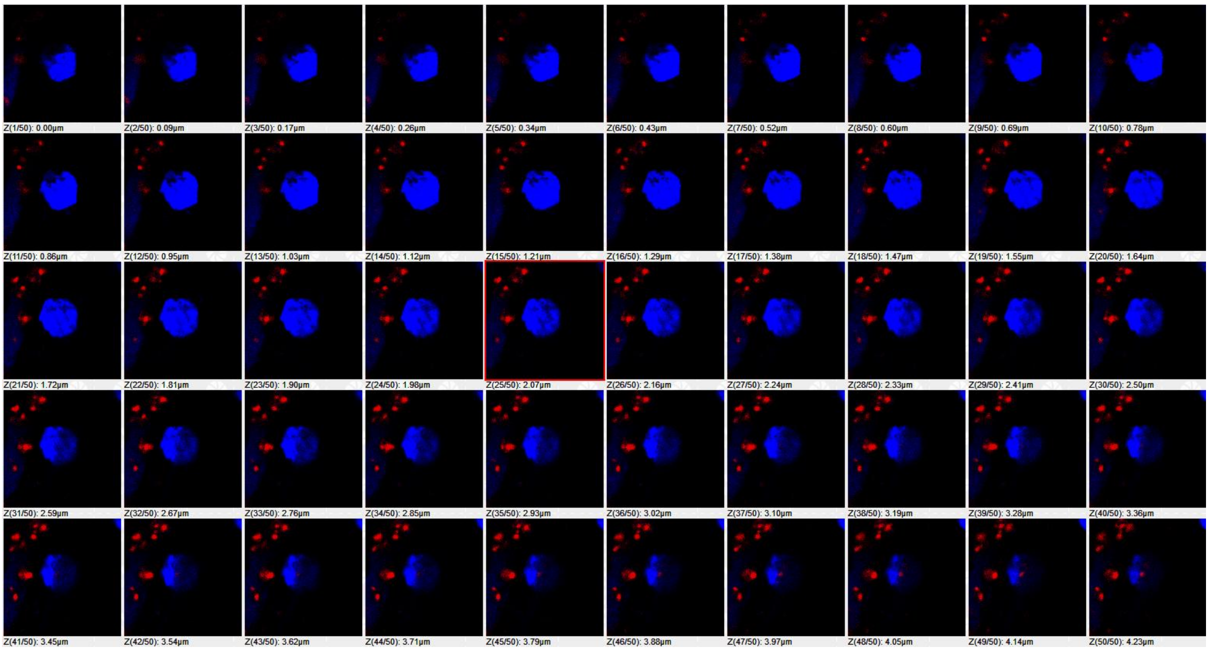


**Figure S9. Confocal z-stack imaging of chloroplast-nucleus organization in lincomycin- grown with anaesthesia-treated leaf tissue.** Representative sequential confocal optical sections (**Z1-Z50**) showing the spatial distribution of chloroplasts and nuclei following combined lincomycin and anaesthesia treatment. Red fluorescence represents chloroplast autofluorescence, whereas blue fluorescence represents nuclear staining. The z-stack demonstrates changes in chloroplast abundance, fluorescence intensity, and spatial organization relative to the nucleus, indicating the combined effects of chloroplast-development inhibition and anaesthesia on leaf cellular organization.

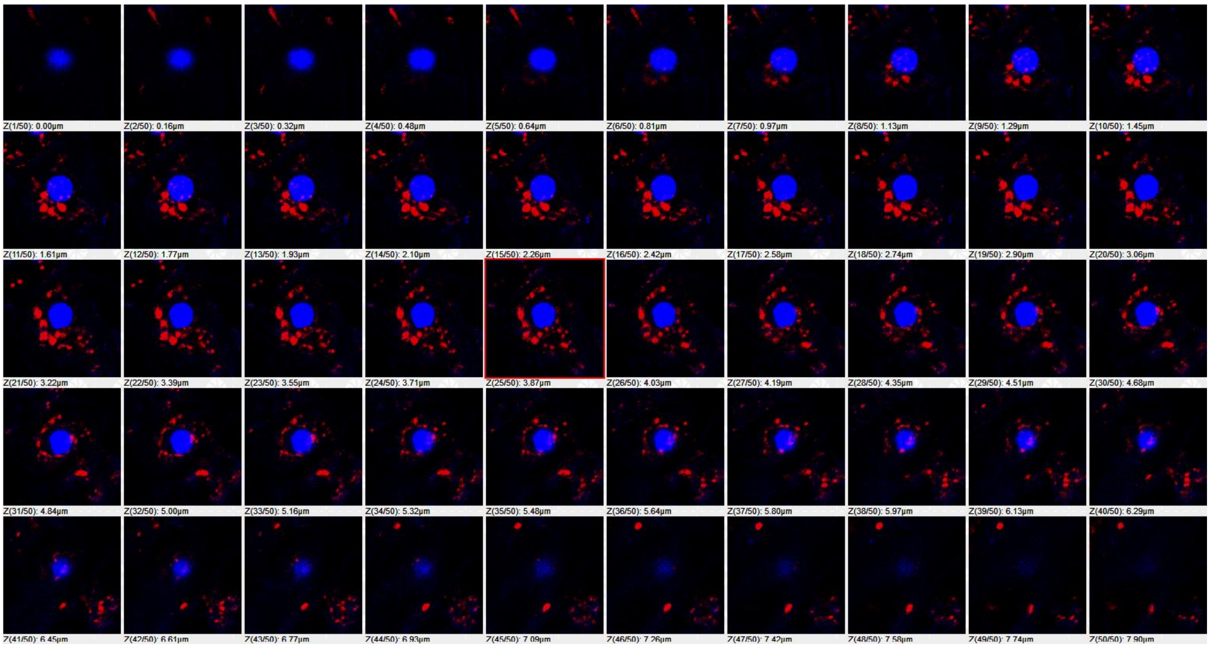


**Figure S10. Confocal z-stack imaging of chloroplast-nucleus organization in stem tissue under control light conditions.** Representative sequential confocal optical sections (**Z1-Z50**) showing the spatial distribution of chloroplasts and nuclei in stem tissue under control light conditions. Red fluorescence represents chloroplast autofluorescence, whereas blue fluorescence represents nuclear staining. The z-stack demonstrates the normal distribution and organization of chloroplasts around the nucleus.

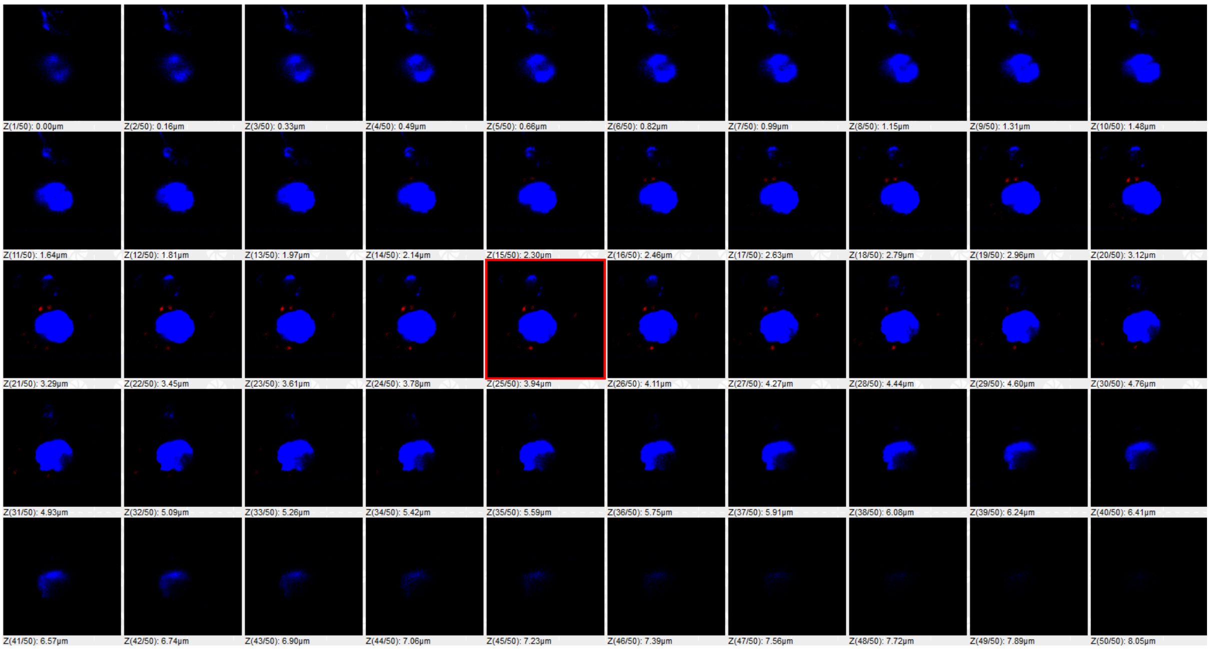

**Figure S11. Confocal z-stack imaging of chloroplast-nucleus organization in dark-grown stem tissue.** Representative sequential confocal optical sections (**Z1-Z50**) showing chloroplast and nuclear organization in stem tissue grown under dark conditions. Red fluorescence represents chloroplast autofluorescence, while blue fluorescence represents nuclear staining. The z-stack reveals reduced chloroplast-associated fluorescence and altered chloroplast distribution, consistent with impaired chloroplast development under darkness.

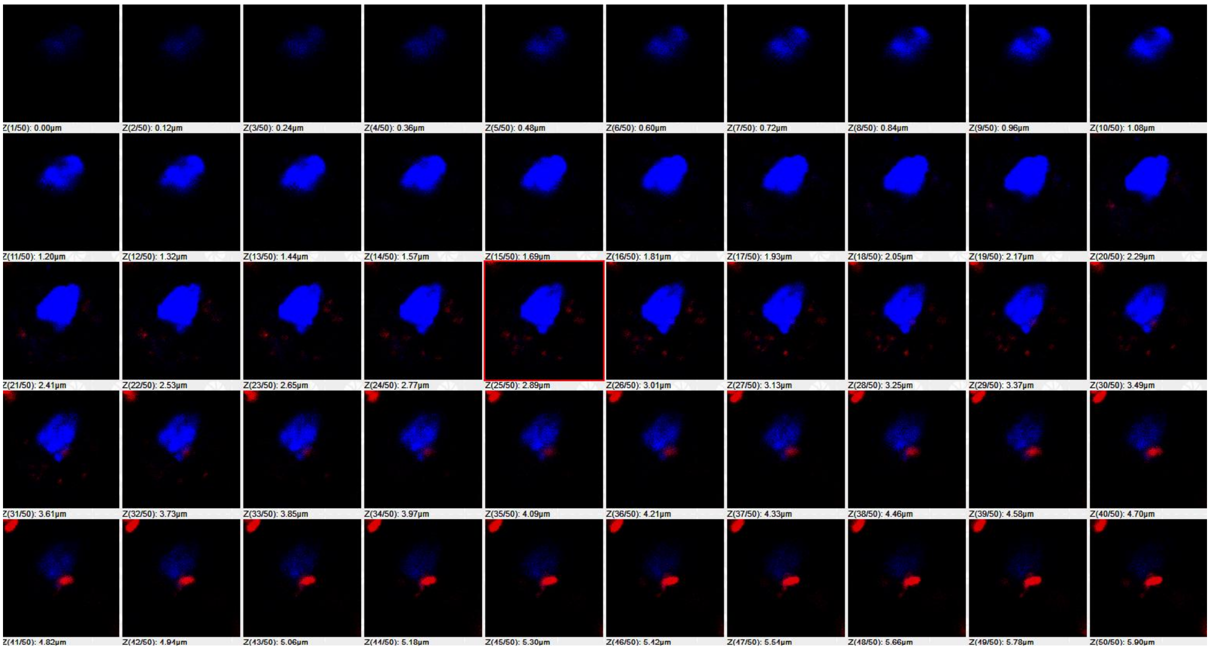


**Figure S12. Confocal z-stack imaging of chloroplast-nucleus organization in lincomycin-grown stem tissue.** Representative sequential confocal optical sections (**Z1-Z50**) showing the spatial distribution of chloroplasts and nuclei following lincomycin treatment. Red fluorescence represents chloroplast autofluorescence, whereas blue fluorescence represents nuclear staining. The z-stack demonstrates reduced and altered chloroplast-associated fluorescence and changes in chloroplast organization relative to the nucleus following inhibition of chloroplast development.

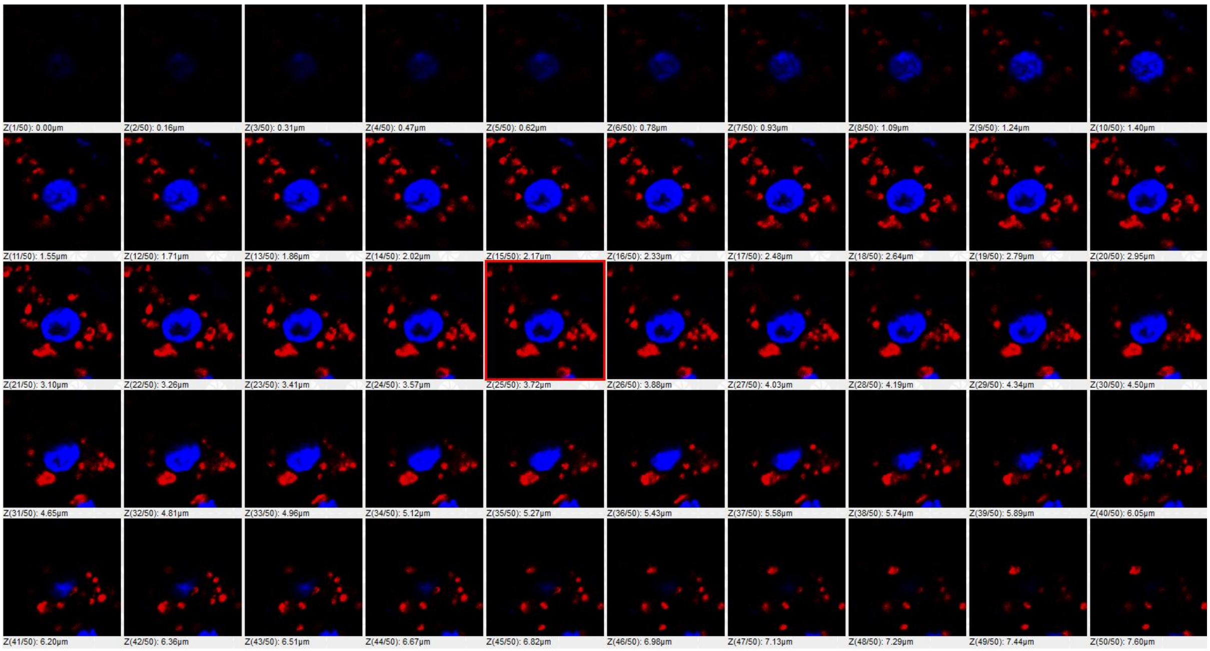


**Figure S13. Confocal z-stack imaging of chloroplast-nucleus organization in anaesthesia-treated stem tissue.** Representative sequential confocal optical sections (**Z1-Z50**) showing chloroplast and nuclear organization following anaesthesia treatment. Red fluorescence represents chloroplast autofluorescence, while blue fluorescence represents nuclear staining. The z-stack illustrates changes in chloroplast distribution, fluorescence intensity, and the z-stack reveals a pronounced peripheral arrangement of nuclear DNA/chromatin following anaesthesia exposure, indicating anaesthesia-associated reorganization of nuclear DNA.

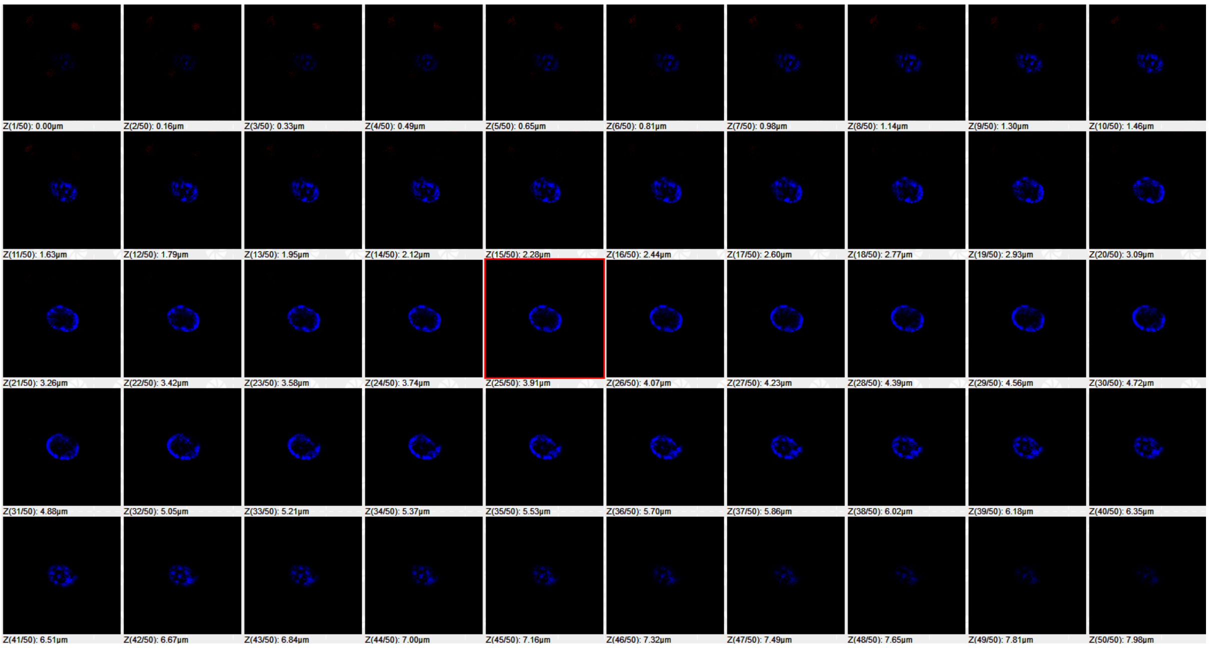


**Figure S14. Confocal z-stack imaging of chloroplast-nucleus organization in dark-grown, anaesthesia-treated stem tissue.** Representative sequential confocal optical sections (**Z1-Z50**) showing the spatial distribution of chloroplasts and nuclei following combined dark growth and anaesthesia treatment. Red fluorescence represents chloroplast autofluorescence, and the z-stack reveals a pronounced peripheral arrangement of nuclear DNA/chromatin following anaesthesia exposure, indicating anaesthesia-associated reorganization of nuclear DNA.

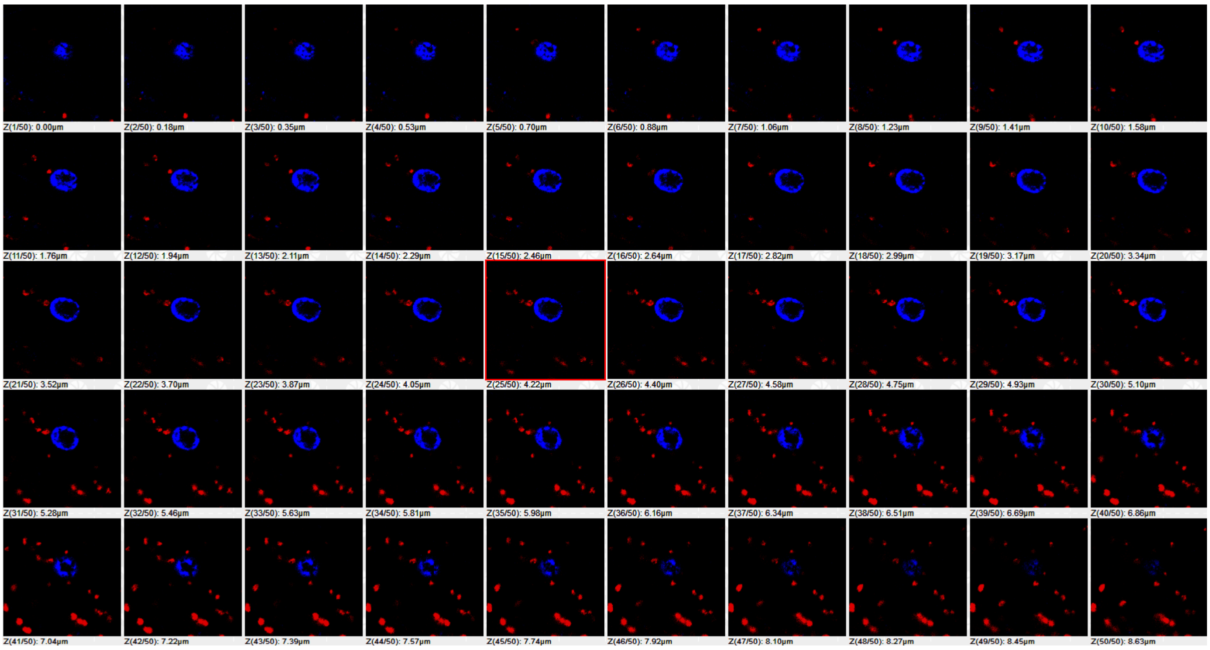


**Figure S15. Confocal z-stack imaging of chloroplast-nucleus organization in lincomycin- grown with anaesthesia-treated stem tissue.** Representative sequential confocal optical sections (**Z1-Z50**) showing chloroplast and nuclear organization following combined lincomycin and anaesthesia treatment. Red fluorescence represents chloroplast autofluorescence, while blue fluorescence represents nuclear staining. The z-stack demonstrates changes in chloroplast-associated fluorescence, abundance, and the z-stack reveals a pronounced peripheral arrangement of nuclear DNA/chromatin following anaesthesia exposure, indicating anaesthesia-associated reorganization of nuclear DNA.

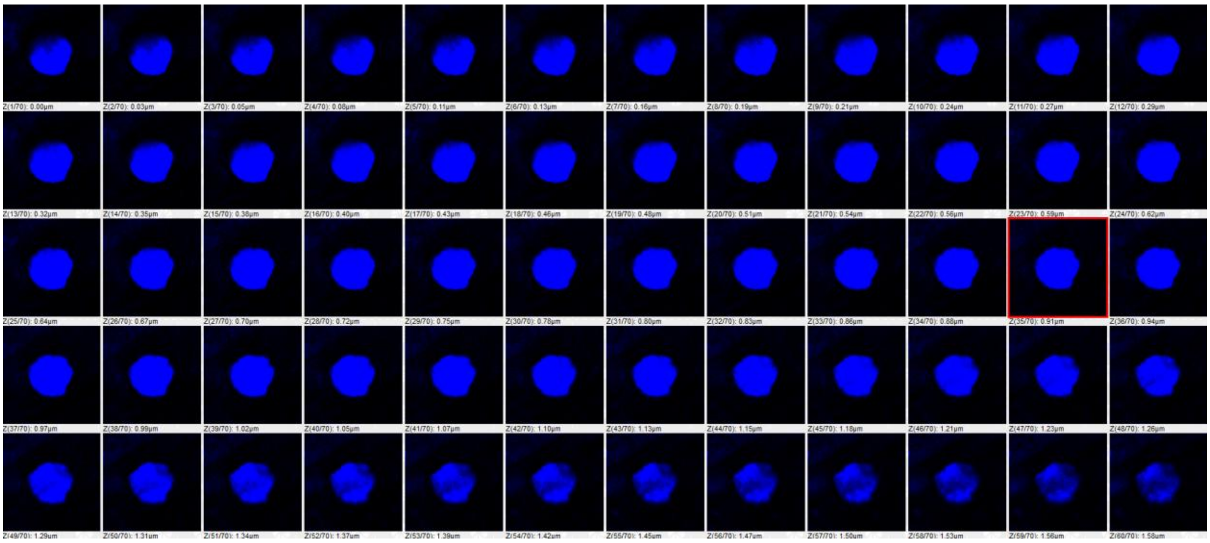

**Figure S16. Confocal z-stack imaging of nuclear organization in root tissue under control light conditions.** Representative sequential confocal optical sections showing nuclear organization in root tissue under control light conditions. Blue fluorescence represents nuclear/DNA staining. The z-stack demonstrates the normal spatial distribution and organization of nuclear DNA within the root cells.

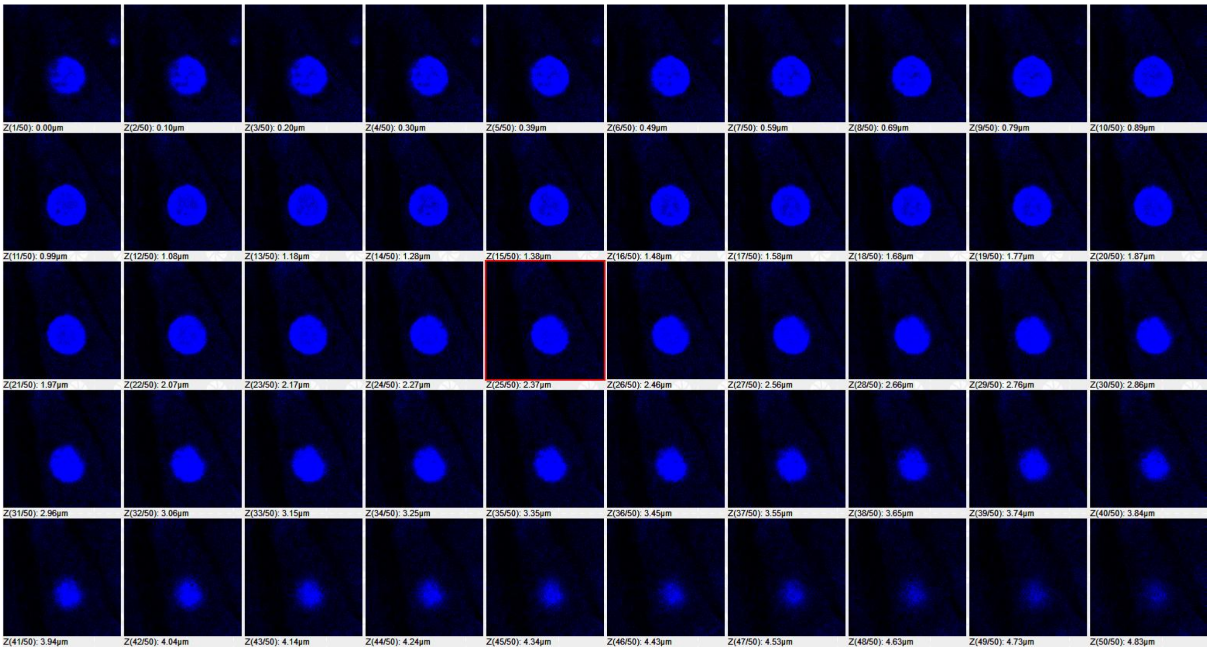


**Figure S17. Confocal z-stack imaging of nuclear organization in dark-grown root tissue.** Representative sequential confocal optical sections showing nuclear organization in root tissue grown under dark conditions. Blue fluorescence represents nuclear/DNA staining. The z-stack demonstrates the spatial organization of nuclear DNA under dark-growth conditions, with changes in nuclear organization compared with the control condition.

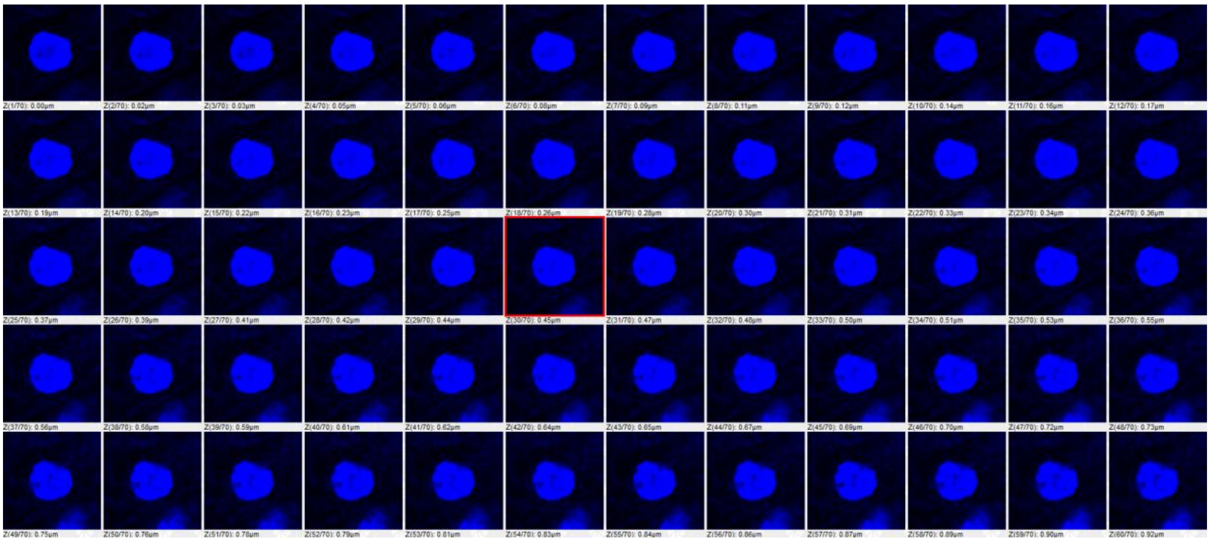

**Figure S18. Confocal z-stack imaging of nucleus organization in lincomycin-grown root tissue.** Representative sequential confocal optical sections showing nuclear organization in root tissue following lincomycin treatment. Blue fluorescence represents nuclear/DNA staining. The z-stack illustrates the spatial distribution of nuclear DNA and provides a comparison of nuclear organization following lincomycin treatment.

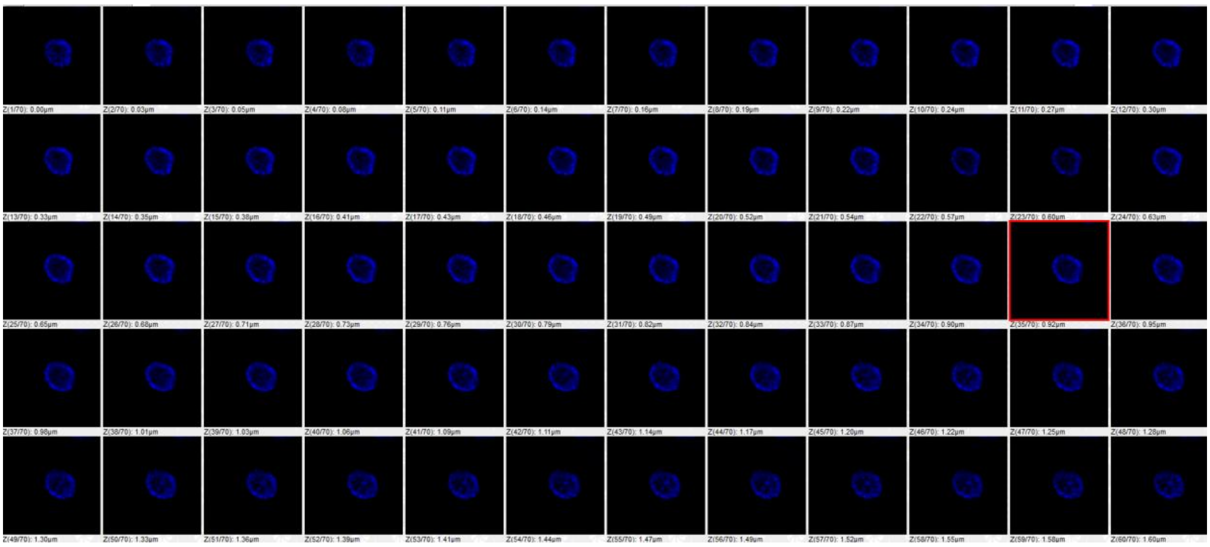

**Figure S19. Confocal z-stack imaging of nuclear DNA organization in anaesthesia-treated root tissue.** Representative sequential confocal optical sections showing nuclear organization following anaesthesia treatment. Blue fluorescence represents nuclear/DNA staining. The z-stack reveals a pronounced peripheral arrangement of DNA/chromatin within the nucleus following anaesthesia exposure, indicating anaesthesia-associated reorganization of nuclear DNA.

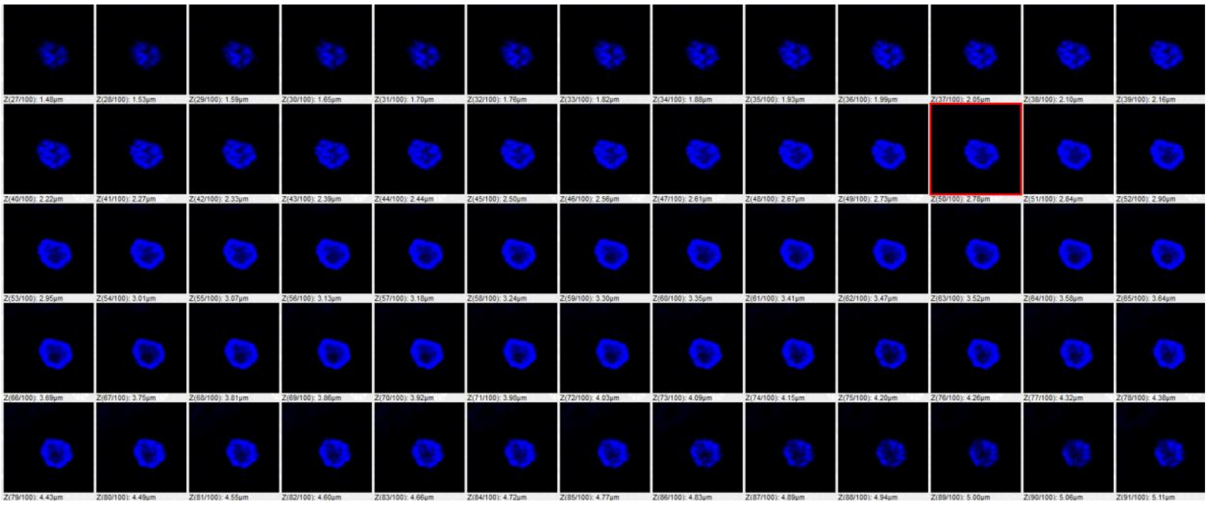

**Figure S20. Confocal z-stack imaging of nuclear DNA organization in dark-grown, anaesthesia-treated root tissue.** Representative sequential confocal optical sections showing nuclear organization following combined dark growth and anaesthesia treatment. Blue fluorescence represents nuclear/DNA staining. The z-stack demonstrates peripheral redistribution of nuclear DNA/chromatin associated with anaesthesia treatment under dark-growth conditions.

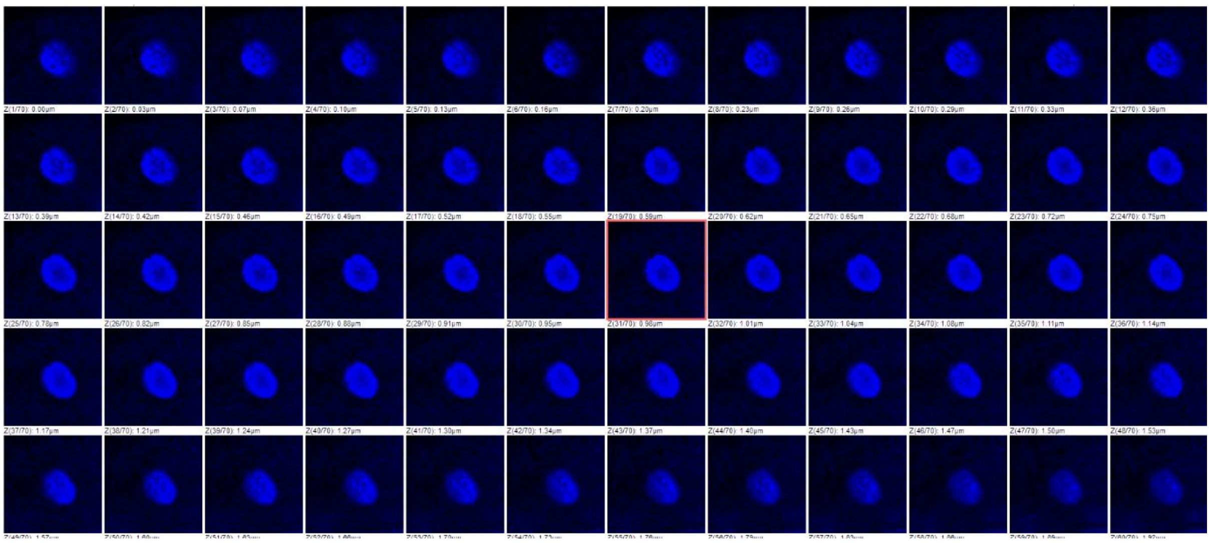

**Figure S21. Confocal z-stack imaging of nuclear DNA organization in lincomycin- grown with anaesthesia-treated root tissue.** Representative sequential confocal optical sections showing nuclear organization following combined lincomycin and anaesthesia treatment. Blue fluorescence represents nuclear/DNA staining. The z-stack demonstrates peripheral organization of nuclear DNA/chromatin, indicating that anaesthesia-associated nuclear reorganization is evident under the combined treatment condition.
